# Single-cell-level digital twins for preterm birth prevention strategies

**DOI:** 10.1101/2025.09.29.679252

**Authors:** Jakob Einhaus, Peter Neidlinger, Olivier Fondeur, Masaki Sato, Alexandra Anronikov, Kotaro Miyazaki, Jonas N. Amar, Kazuo Ando, Valentin Badea, Dyani K. Gaudilliere, Maximilian Sabayev, Dorien Feyaerts, Maïgane Diop, Amy S. Tsai, Amelie Cambriel, Edward A. Ganio, Romain Lagarde, Emmeline O’Kelly, Ina A. Stelzer, Julien Hedou, Ronald J. Wong, Yair J. Blumenfeld, Deirdre J. Lyell, Gary M. Shaw, Tomiko T. Oskotsky, Marina Sirota, Linda Giudice, David K. Stevenson, Nima Aghaeepour, Brice Gaudilliere

## Abstract

Digital twin models can accelerate therapeutic development by enabling low-risk testing of candidate interventions. In preterm labor (PTL), a major pregnancy complication where clinical trials face unique ethical and financial barriers, digital twins are especially valuable for evaluating new therapies targeting immune dysfunctions driving PTL. Yet, current models lack single-cell resolution, limiting detection of cell-type-specific mechanisms, off-target effects, and the design of personalized interventions. We present Simulated Immunome Modeling of Clinical Outcomes (SIMCO), a single-cell-level digital twin framework that models immunomodulatory treatment effects on the timing of labor using immunome-wide, single-cell simulations. SIMCO’s digital twins are trained and validated on a newly generated mass cytometry atlas of the pregnant immunome exposed to nine candidate drugs preselected for PTL prevention. Applying SIMCO to an independent cohort of pregnant individuals, we simulate treatment effects on gestational length, screening for candidate drugs that delay labor timing and providing system-level mechanistic insight for each drug candidate. Tetrahydrofolate, maprotiline, and the combination of aspirin and lansoprazole emerged as top candidates for PTL prevention, delaying labor onset primarily through enhanced mTOR signaling in innate immune cells and attenuated JAK/STAT signaling in naïve CD4⁺ T cells. The codebase is available at https://github.com/ofondeur/SIMCO/.

## INTRODUCTION

Preterm birth, defined as birth before 37 completed weeks of gestation, remains the leading cause of child mortality under age five worldwide.^1^ One in ten pregnancies ends preterm, leading to significant neonatal complications. Most cases (60–70%) are spontaneous preterm labor (sPTL) and can occur without warning signs or known risk factors.^2^ While multiple factors contribute to sPTL, increasing evidence points to dysregulation of innate and adaptive immune adaptations that maintain pregnancy from conception to labor.^3–11^ Early pregnancy is dominated by tolerogenic mechanisms, such as the expansion of regulatory T cells (Tregs), which promote maternal–fetal tolerance.^12–15^ Late pregnancy involves activation of pro-inflammatory pathways in neutrophils, monocytes, macrophages, and natural killer (NK) cell subsets.^16–18^ These temporal dynamics culminate in an inflammatory cascade of parturition.^19–21^ In sPTL, anticipated immune adaptations of healthy pregnancies occur prematurely or inappropriately.^22^

Understanding the differences in gestational immune dynamics between pregnancies resulting in sPTL and those delivering at term is critical for identifying immunomodulatory interventions that could influence timing of labor onset.^23,24^ Current sPTL preventive strategies, including low-dose aspirin^25^ and progesterone,^26,27^ have shown modest and inconsistent results in clinical trials, likely benefiting some but not all patients.^28^ Given the ethical and financial constraints on clinical trials during pregnancy, immune digital twins offer a transformative approach to accelerate the development of immunomodulatory drug candidates for sPTL.^29,30^ Incorporating single-cell resolution can equip digital twin models to capture immune system–wide drug effects, reveal cell-type–specific mechanisms, identify off-target responses, and guide the rational design of personalized interventions.

Mass cytometry enables single-cell immune profiling with high-dimensional functional readouts including cell-surface receptors, cytokines, and phosphorylation states.^31^ Beyond characterizing immune states, the rich datasets generated by mass cytometry can be harnessed by machine learning to extract biomarkers of clinical outcomes. For example, the recently developed Stabl algorithm uses artificial noise to select informative features and build sparse, interpretable prediction models.^32^ Applied to peripheral blood samples across gestation, mass cytometry combined with Stabl revealed conserved temporal trajectories of immune cell signaling activity, enabling prediction of labor timing in both term and preterm pregnancies.^5,33^ Besides biomarker discovery, mass cytometry can also serve as a high-throughput screening platform to study cell-type and pathway specificity of immunomodulatory therapeutics.^34^ With recent advances in computational modeling, these observed drug effects can then be parsed at the single-cell level. In particular, optimal transport (OT) has emerged as a powerful mathematical framework to model single-cell immune perturbation responses.^35^ For mapping unpaired distributions with and without perturbation, CellOT by Bunne et al. leverages a neural network for efficient OT solving.^36^ Its robust performance on unseen data enables personalized treatment response modeling beyond the original perturbation dataset.

In this study, we present Simulated Immunome Modeling of Clinical Outcomes (SIMCO), a single-cell-level digital twin framework that models immunomodulatory treatment effects on clinical outcomes by integrating simulated immune responses with multivariable modeling. Trained on a new single-cell mass cytometry atlas of the pregnant immunome, SIMCO’s digital twins predict patient-specific immune modulation and resulting effects on gestational length for nine FDA-approved, low-risk (category A/B) drug candidates – eight single agents and one combination – preselected through a published computational drug repurposing pipeline for their potential to prevent preterm birth.^23^

## RESULTS

### Simulating treatment effects from single-cell-level digital twins

Simulating the therapeutic potential of immunomodulatory treatment requires digital twins of the immunome that not only predict clinical outcomes but also forecast how specific treatments would alter these outcomes in individual patients. SIMCO addresses this need by integrating simulation of single-cell responses with clinical outcome modeling (**Figure 1, Methods, and Supplementary Figure S1**):

1. First, SIMCO is trained on high-dimensional single-cell perturbation data (e.g., flow and mass cytometry, single-cell transcriptomics) to encode drug-perturbed immune responses at single-cell resolution using OT learning with CellOT. CellOT leverages an input-convex neural network that is highly accurate in predicting heterogeneous single-cell perturbation responses.^36^
2. Second, SIMCO projects the learned single-cell behavior onto a separate immunome dataset, comprising only baseline samples (unstimulated and untreated) with linked clinical outcomes. From these baseline samples, SIMCO simulates single-cell responses to perturbations (e.g., ex vivo stimulation or drug treatment). This simulation generates digital twins of each patient’s immunome with and without exposure to the candidate drugs.
3. Third, SIMCO builds a multivariable predictive model of the clinical outcome with Stabl, a sparse machine learning algorithm, on a training set of patients using the simulated immune data. In a held-out test set of patients, SIMCO compares the predictions for each patient’s untreated and treated digital twin. This difference quantifies the predicted drug effect on the outcome, enabling SIMCO to model the immunomodulatory potential of candidate drugs and to personalize treatment strategies.

**Figure 1:**
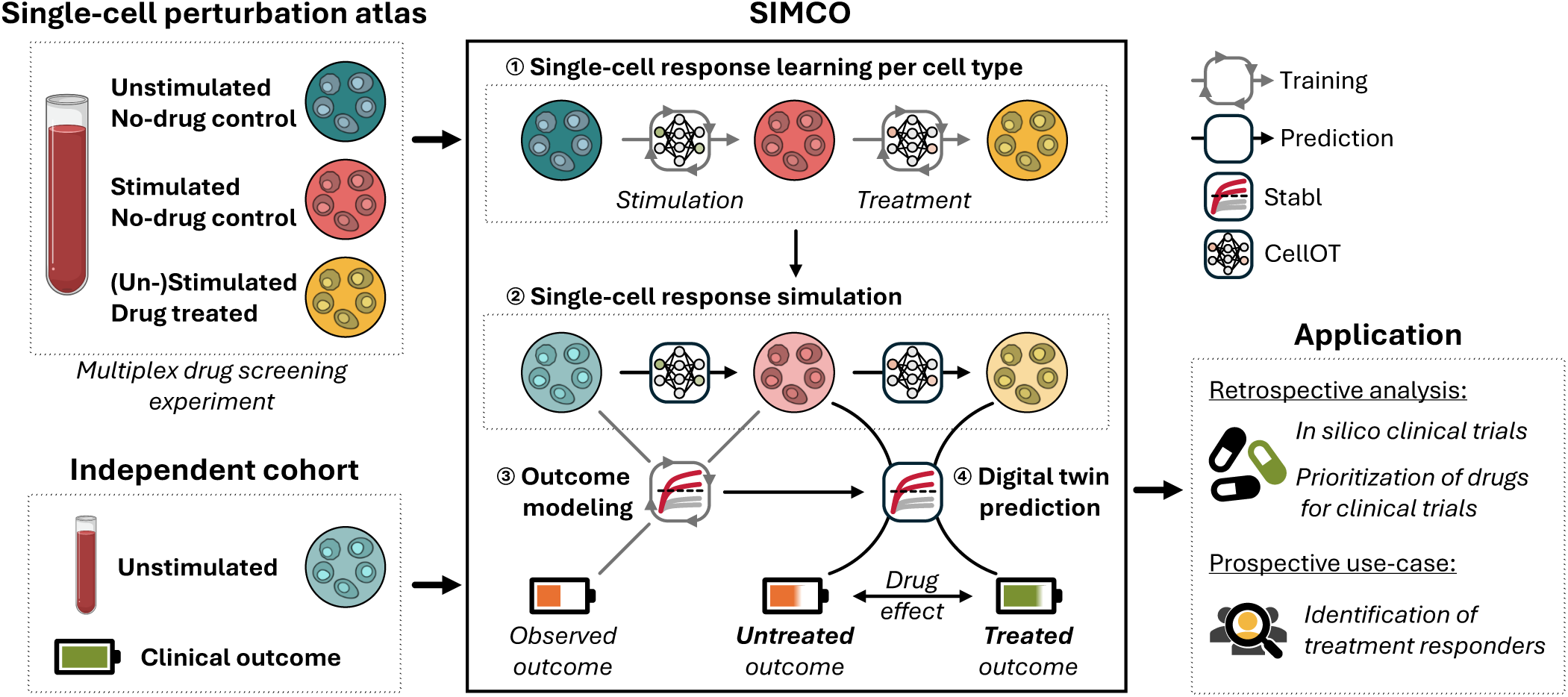
Overview over the single-cell-level digital twin framework SIMCO. Simulated Immunome Modeling of Clinical Outcomes (SIMCO) is a single-cell-level digital twin framework that combines simulated immune responses with multivariable modeling of clinical outcomes to evaluate the immunomodulatory potential of candidate drugs. In vitro perturbation experiments (i.e., analysis of ex vivo stimulation responses to inflammatory ligands in the presence or absence of drug candidates) are performed to generate a single-cell perturbation atlas of immunomodulatory treatment effects. SIMCO learns optimal transport maps (using CellOT) that encode the observed single-cell perturbation responses. For baseline samples from an independent clinical cohort linked to a clinical outcome, SIMCO simulates single-cell responses by applying the learned optimal transport maps. A sparse machine learning model (Stabl) is trained on the simulated single-cell data before treatment (light turquoise, light red) to model the observed outcome. Applied to an unseen test set of patients, SIMCO provides a readout of simulated drug effects on the outcome as the difference in outcome prediction between the treated and untreated digital twin (light red, light yellow). SIMCO-generated immune digital twins can be used for in silico clinical trials and prioritization of candidate drugs for clinical trial testing, as well as for personalization of treatment strategies.

### A single-cell perturbation atlas capturing pregnancy-relevant immune profiles

To generate a mass cytometry single-cell perturbation atlas, we enrolled eight non-pregnant and 11 pregnant volunteers for a multiplexed drug screen assay measuring immunomodulatory effects of nine candidate drugs for the prevention of sPTL (**Figure 2a**, **Table 1**). A second trimester sampling timepoint was chosen to capture the influence of candidate drugs on the pregnant immunome during the maintenance phase of gestation and before proinflammatory pathways of parturition become increasingly activated. Non-pregnant controls were sampled in the early follicular phase of their menstrual cycle to capture a low-hormone state. The candidate drugs – chlorthalidone, maprotiline, metformin, lansoprazole, pravastatin, rifabutin, salicylic acid (SA), salicylic acid + lansoprazole (SALPZ), and 5-methyltetrahydrofolate (THF) – were preselected using a previously published computational drug repurposing pipeline based on their potential to positively influence sPTL-associated gene signatures.^23^ Only candidate drugs considered relatively safe for use in pregnancy (FDA category A/B) that can be administered orally were included in the assay. For each candidate, we used the compound form predominantly present in the bloodstream after oral administration, which in some cases corresponds to an active metabolite (e.g., THF, SA). Experimental concentrations were selected to reflect in vivo plasma levels following standard oral dosing (**Table 2**).

**Figure 2:**
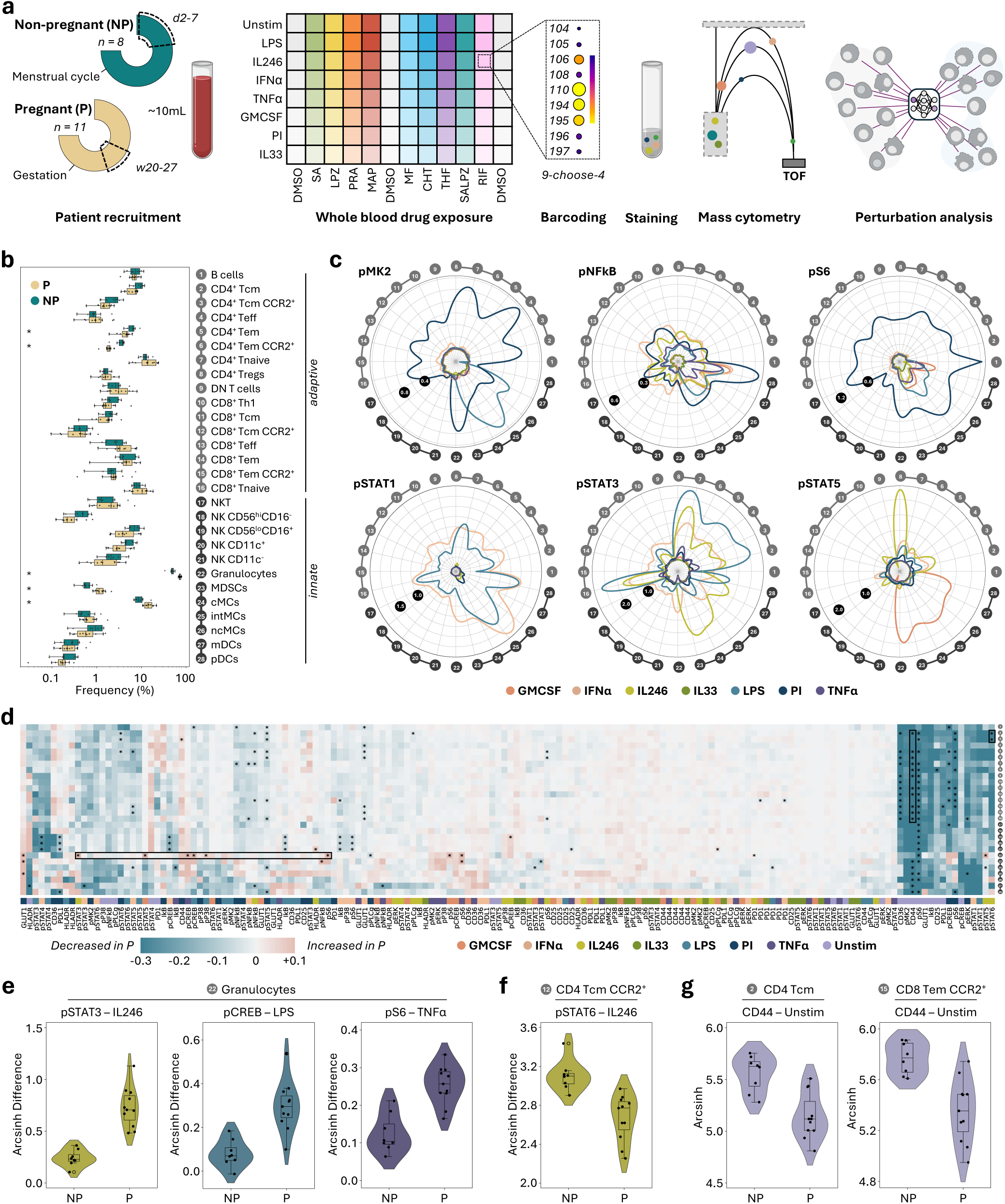
A single-cell immune perturbation atlas of the second-trimester pregnancy immunome. **a.** In 19 individuals, including eight non-pregnant (NP) between days two to seven of the menstrual cycle and 11 pregnant (P) between 20 to 27 weeks of gestation, immunomodulatory effects of nine candidate drugs (eight single drugs and one drug combination) were screened in a 96-plex ex vivo whole blood immunoassay. **b.** Proportional abundance of 28 cell populations in pregnant and non-pregnant individuals. Percent of all live leukocytes (granulocytes) and percent of mononuclear cells (all other cell types) are depicted. Significance (p < 0.05, Mann-Whitney U test, Benjamini-Hochberg corrected) is indicated. **c.** Radial plots show absolute arcsinh difference between stimulated and unstimulated samples across 28 immune cell populations for selected functional markers. **d.** Heatmap of differences in average immune responses to stimulation between pregnant and non-pregnant individuals. Stars indicate significant differences (p < 0.05, Wilcoxon rank-sum test, Benjamini-Hochberg corrected across all features with a group difference > 0.05 arcsinh). Feature groups with significant differences (individual features plotted in e-g) are highlighted with boxes. e-g. Example features of significant differences between pregnant and non-pregnant individuals. Pregnant individuals showed upregulated immune responses in innate immunity (granulocytes, e), downregulated adaptive immune responses (CD4+ central memory T cells, f), and downregulated CD44 expression (**g**). Depicted is arcsinh difference to unstimulated (for stimulation responses) or arcsinh-transformed signal (for unstimulated). Candidate drugs are abbreviated as follows: chlorthalidone (CHT), maprotiline (MAP), metformin (MF), lansoprazole (LPZ), pravastatin (PRA), rifabutin (RIF), salicylic acid (SA), salicylic acid + lansoprazole (SALPZ), and 5-methyltetrahydrofolate (THF). Dimethyl sulfoxide (DMSO) was used as no-drug control.

**Table 1:** Clinical information and patient demographics for pregnant and non-pregnant volunteers. Percentage and interquartile range (IQR) within each cohort are indicated.

|  | Non-pregnant individuals (n = 8) | Pregnant individuals (n = 11) |
| --- | --- | --- |
| Age (years) | 32 (29-35, IQR) | 30 (29-32, IQR) |
| Race |  |  |
| White | 5 (62.5%) | 4 (36.4%) |
| Asian | 2 (25.0%) | 4 (36.4%) |
| Black | 1 (12.5%) | 0 |
| Other | 0 | 3 (27.3%) |
| Ethnicity |  |  |
| Non-Hispanic | 8 | 8 |
| Hispanic | 0 | 0 |
| Other | 0 | 3 |
| Pregnancy information |  |  |
| GA at sampling time |  | 22.4 (20.9 - 22.8, IQR) |
| Pre-pregnancy BMI |  | 24.6 (22.4 - 27.0, IQR) |
| GA at delivery |  | 40.0 (39.5-40.05, IQR) |
| Medication |  |  |
| Prenatal vitamin |  | 7 (63.6%) |
| Omeprazole prenatal |  | 1 (9.1%) |
| Contraception |  |  |
| Oral contraception | 1 (9.1%) |  |
| Copper IUD | 1 (9.1%) |  |
| Hormonal IUD | 2 (18.2%) |  |
| Cycle day | 3 (3-4, IQR) |  |
| Medical history |  |  |
| Hyperlipidemia |  | 1 (9.1%) |
| Factor V Leiden |  | 1 (9.1%) |
| PCOS | 1 (12.5%) | 1 (9.1%) |
| IUI |  | 1 (9.1%) |

**Table 2:** Candidate drugs for the prevention of spontaneous preterm labor. The screened compound, the used assay concentration, and the abbreviation (Abb.) of the compound in the figures are listed.

| Drug candidate | Clinical reference dose | Cmax (ng/mL) | T1/2 (h) | Relevant metabolites | Assay concentration (ng/mL) | Screened compound | Abb. | References |
| --- | --- | --- | --- | --- | --- | --- | --- | --- |
| Chlorthalidone | 12.5 mg | 75 | 32 | None | 75 | Chlorthalidone | CHT | <sup>37</sup> |
| Maprotiline | 75 mg | 24 | 40 | Desmethyl-maprotiline | 10 | Maprotiline | MAP | <sup>38</sup> |
| Metformin | 500 mg | 1100 | 4 | None | 800 | Metformin | MF | <sup>39</sup> |
| Lansoprazole | 30 mg | 1000 | 2 | 5'-hydroxy-lansoprazole | 400 | Lansoprazole | LPZ | <sup>40</sup> |
| Pravastatin | 10 mg | 15 | 2.5 | 3'α-isopravastatin | 7.5 | Pravastatin | PRA | <sup>41</sup> |
| Rifabutin | 150 mg | 200 | 36 | 25-O-deacetyl-rifabutin | 10 | Rifabutin | RIF | <sup>42</sup> |
| Aspirin | 81 mg | 5700 | 2.5 | Salicylic acid | 2500 | Salicylic acid | SA | <sup>43</sup> |
| Folic acid | 800 µg | 15 | 5.5 | 5-Methyl-THF | 75 | 5-Methyl-THF | THF | <sup>44</sup> |

Each screening experiment consisted of 96 conditions requiring only 100 µL of whole blood per condition. A two-hour, whole blood stimulation allowed us to examine physiologically relevant cell type-and stimulation-specific responses. Functional responsiveness of 28 immune cell populations was assessed as expression changes in 12 intracellular phospho-proteins and eight activation markers (**Supplementary Figure S2, Supplementary Table S1**). All conditions were analyzed simultaneously using mass cytometry by leveraging a newly developed nine-choose-four doublet-removing mass-tag barcoding scheme (see **Supplementary Figure S3 and Methods**). In total, our single-cell perturbation atlas comprised 218 million leukocytes across 80 perturbation scenarios.

In the absence of drug treatment (0.5 % dimethyl sulfoxide, DMSO-control), pregnant and non-pregnant individuals exhibited distinct immune profiles. Several innate immune populations – such as granulocytes, classical monocytes (cMCs), and myeloid-derived suppressor cells – were proportionally expanded during mid-pregnancy, whereas CD4⁺ effector memory T cells were reduced (**Figure 2b**). Functional responses to seven stimuli were quantified for each cell type, revealing highly specific response profiles for different signaling pathways induced to each stimulation condition (**Figure 2c)**. Out of 4,480 immune activation features, 163 differed significantly between the two groups (**Figure 2d**).

Specifically, in samples from pregnant compared to non-pregnant participants, we observed increased innate immune signaling in granulocytes in response to interleukin (IL)-2/4/6, lipopolysaccharide (LPS), and tumor necrosis factor (TNF)α (**Figure 2e**), suppressed adaptive immune responses, e.g., in pSTAT6 in CD4^+^ central memory T cells in response to IL-2/4/6 (**Figure 2f**), and reduced expression of CD44 on innate and adaptive immune cells in unstimulated samples (**Figure 2g**). These findings highlight the global adaptations of the maternal peripheral immune system during pregnancy and recapitulate important innate and adaptive cell responses previously implicated in the maintenance of healthy pregnancies.^4,33,45^

### Candidate drugs show pregnancy-specific immunomodulatory properties

From the single-cell perturbation atlas, we characterized each candidate drug’s immunomodulatory profile when administered in the context of the pregnant immunome. For each candidate drug, we analyzed 4,480 immune cell features comparing the drug-treated measurements to their respective DMSO controls. This provided an in-depth analysis of cell type-and pathway-specific concerted effects that each candidate drug exerted on the immune cells’ responses and interactions occurring over the two-hour stimulation period (**Figure 3a, Supplementary Figure S4**). For example, SA globally enhanced JAK-STAT responses in response to LPS stimulation, while differentially regulating these pathways under PMA/ionomycin (PI) stimulation in a highly cell type-specific manner. As such, SA increased the pSTAT5 activation in CD4^+^ Tregs after PI stimulation but decreased pSTAT5 in CD8^+^ T cell subsets.

**Figure 3:**
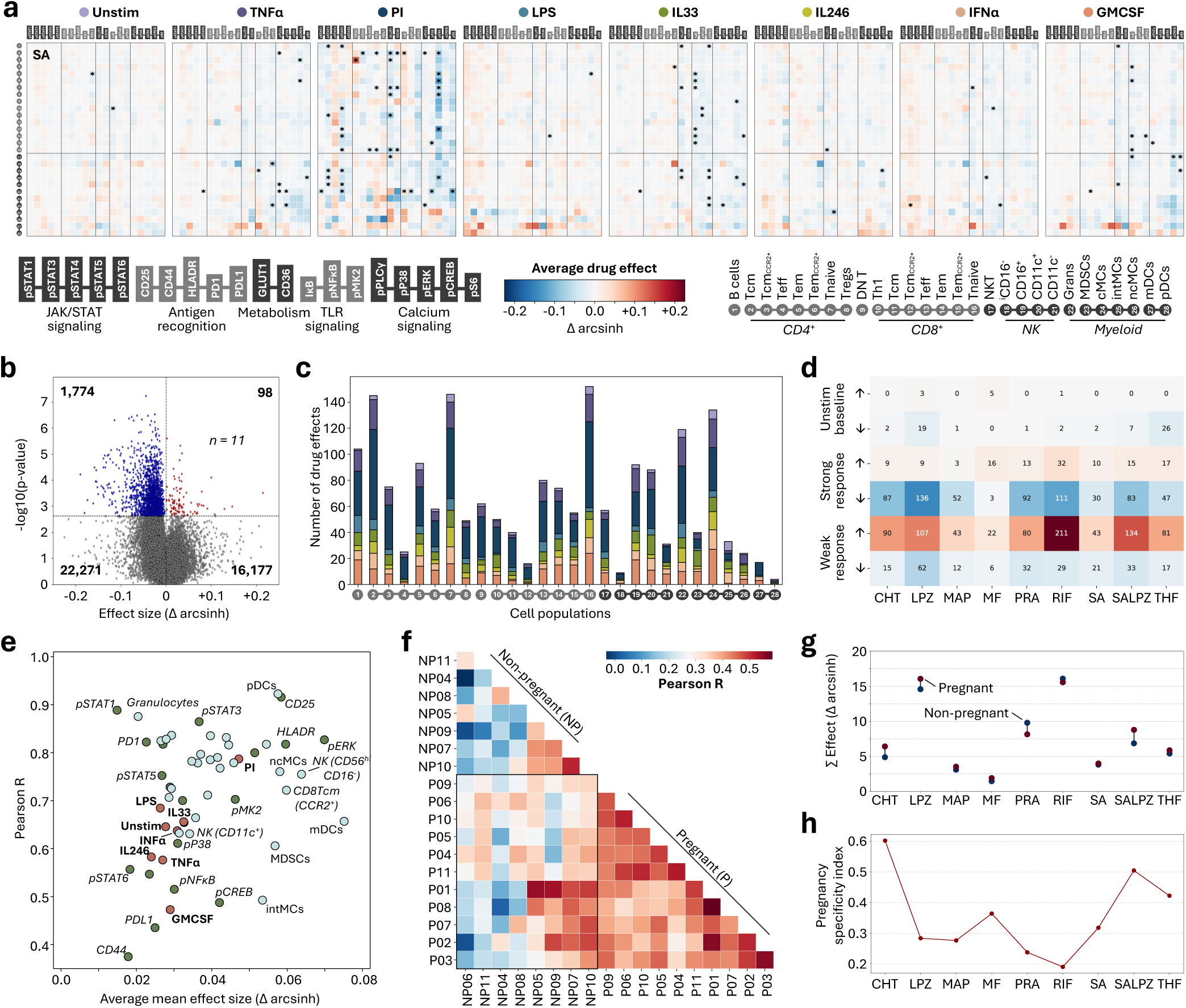
Candidate drugs show pregnancy-specific immunomodulatory properties. **a.** Heatmap of drug effects exerted by SA on the pregnant immunome (n = 11). The arcsinh difference between treated and untreated is plotted and significant effects (see b) are labeled by stars. **b.** Volcano plot of all 40,320 drug effects for pregnant individuals (n = 11). Mean drug effect size and univariate p-value (paired t-test) are plotted. Significant drug effects with a p-value < 0.05 after Benjamini-Hochberg correction are colored in blue (decreasing) or red (increasing). **c.** Number of significant drug effects (see b)per cell population, colored by stimulation condition. **d.** Number of significant drug effects (see b) by candidate drug, categorized as stimulatory/ amplifying (↑) or inhibiting/neutralizing (↓) effects affecting either unstimulated (baseline) immune activity, strong stimulation responses (> 0.1 arcsinh difference between unstimulated and stimulated), or weak stimulation responses (< 0.1 arcsinh difference). **e.** Cross-cohort correlation analysis between cohort-level averages of pregnant (P) and non-pregnant (NP) for significant drug effects (see **b**) grouped by protein (green), cell type (light blue), or stimulation (orange) targeted. The average Pearson R is plotted against the average effect size. **f.** Correlation matrix of all significant drug effects (see b) between individuals. **g.** Average cumulative effect size for all significant drug effects (see b) comparing pregnant and non-pregnant. **h.** Average pregnancy specificity index of all significant drug effects (see b). Candidate drugs are abbreviated as follows: chlorthalidone (CHT), maprotiline (MAP), metformin (MF), lansoprazole (LPZ), pravastatin (PRA), rifabutin (RIF), salicylic acid (SA), salicylic acid + lansoprazole (SALPZ), and 5-methyltetrahydrofolate (THF).

Across all treatments, we identified 1,872 significant drug effects (paired t-test, *p* < 0.05, Benjamini-Hochberg corrected; **Figure 3b**). Most drug effects (94.8%) were inhibitory. Naïve CD4^+^ and CD8^+^ cells as well as central memory CD4^+^ T cells were the most frequently targeted immune cell populations, followed by granulocytes and cMCs among innate cell types (**Figure 3c**). PI and granulocyte-macrophage colony-stimulating factor (GMCSF) stimulations were most frequently modulated.

We categorized drug effects in the following groups: inhibitory/neutralizing or stimulatory/ amplifying treatment effects affecting 1) baseline immune cell activity, 2) strong stimulation responses (arcsinh difference > 0.1 between unstimulated and stimulated), or 3) weak stimulation responses (arcsinh difference < 0.1; **Figure 3d**). Most effects amplified weak or inhibited strong stimulation responses. Overall, all tested treatments showed cell type-and pathway-specific immunomodulatory profiles that may be leveraged to counteract disease-associated immune dysregulation.

We next investigated whether the identified treatment effects were pregnancy-specific. For each protein marker, stimulation, and cell type, we correlated cohort-level average drug effects between the pregnant and non-pregnant cohorts. This cross-cohort correlation analysis revealed conserved treatment effects on certain protein markers (e.g., pSTAT1, pSTAT3, and CD25), stimulation conditions (e.g., PI), and cell types (e.g., granulocytes and plasmacytoid dendritic cells) across both cohorts (**Figure 3e**). In contrast, effects involving PD-L1, CD44, or GMCSF stimulation differed between pregnant and non-pregnant individuals. Correlating individual-level drug effects between individuals showed that the achieved drug effects were more consistent among pregnant individuals (Pearson R range = 0.3 – 0.5) than among non-pregnant individuals (Pearson R range = 0 – 0.5), reflecting lower variability in treatment responses in the pregnant group (**Figure 3f**). While the cumulative effect size per drug was generally comparable between cohorts, lansoprazole and SALPZ exhibited stronger cumulative effects in pregnant individuals, whereas pravastatin showed a reduced effect (**Figure 3g**). We calculated a pregnancy-specificity index for each drug-feature combination, ranging from 0 (equal effect in both groups) to 1 (effect present only in pregnant individuals, see **Methods**). On average, pregnancy-specificity was highest for effects mediated by SALPZ and chlorthalidone (**Figure 3h**).

Notably, the combination therapy SALPZ achieved a unique immunomodulatory profile compared to its individual components, SA and lansoprazole (**Supplementary Figure S5**). SA alone increased JAK-STAT signaling in response to LPS, whereas lansoprazole and SALPZ exerted inhibitory effects on the same pathway. Lansoprazole strongly suppressed calcium signaling in response to PI stimulation, an effect that was less pronounced with SALPZ and absent with SA. In response to IL-2/4/6, lansoprazole enhanced pSTAT3 and pSTAT6 activation, while SALPZ attenuated these signals. To assess the interaction between the two drugs, we compared the observed effects of SALPZ to the summed effects of SA and lansoprazole, classifying them as synergistic, additive, or antagonistic effects. Across all features, SALPZ effects were synergistic in 3%, additive in 79%, and antagonistic in 18%. Most synergistic interactions involved JAK-STAT-signaling following LPS or IFNα stimulation.

### SIMCO accurately simulates cell type-specific perturbation responses

Perturbation modeling approaches at the single-cell level can leverage millions of data points to provide accurate predictions across heterogeneous individuals. SIMCO uses the optimal transport algorithm CellOT to encode the observed perturbation responses as generalizable transport maps, under the hypothesis that the multivariable marker profile of each cell prior to stimulation or treatment is predictive of its response. For each cell type, SIMCO was trained to model 1) stimulation responses from DMSO-control unstimulated samples and 2) drug-treatment responses within each stimulation condition from DMSO-control samples. The trained models included only the 13 functional markers that overlapped between the perturbation dataset and the clinical dataset from Stelzer et al. (**Figure 4a**).

**Figure 4:**
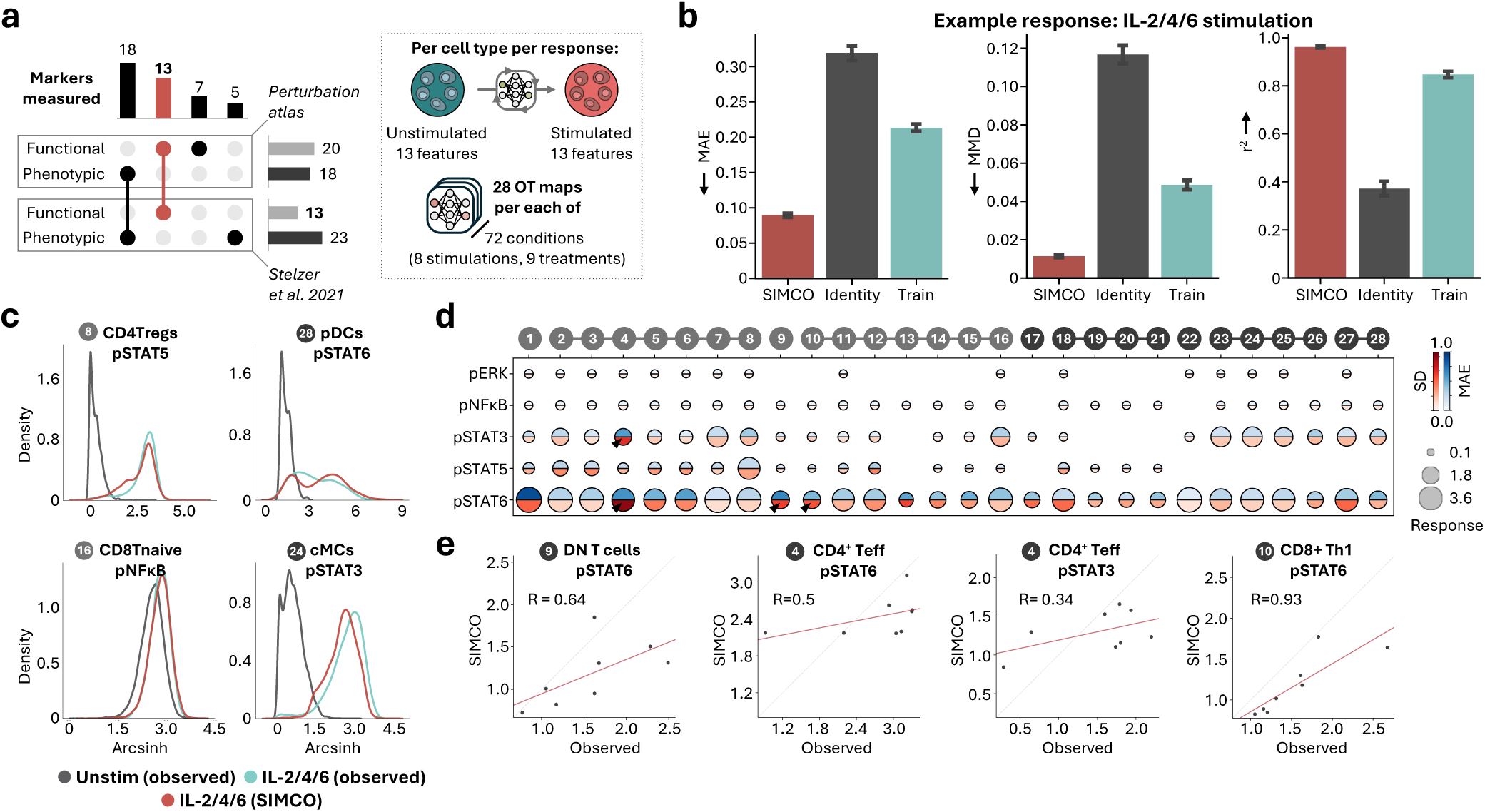
Comparing observed and simulated immune responses to validate SIMCO’s single-cell simulation performance. **a.** Panel alignment between the generated single-cell perturbation atlas and the independent mass cytometry data from Stelzer et al. 2021. In total, 13 functional protein measurements overlapped between the two data sets. Separate OT maps were learned per cell type and response, each predicting 13 functional protein marker changes. The cell types of the two studies were matched, using the same gating strategy based on the 18 overlapping phenotypic markers. **b.** To evaluate SIMCO’s single-cell simulation performance, we compared it to identity and training mean performance in 4-fold cross-validation within the single-cell perturbation atlas. Mean absolute error (MAE), maximum mean discrepancy (MMD), and R2 for the predicted population medians are shown for IL-2/4/6 stimulation responses across all cell types (364 features). **c.** Distributions of protein expression at baseline and in response to IL-2/4/6, comparing observed and SIMCO-simulated single-cell distributions for selected features. **d.** Heatmap of the MAE and the observed standard deviation (SD) for features with an IL-2/4/6 response greater than 0.1 arcsinh difference. **e.** Correlation (Spearman R) between SIMCO-simulated and observed feature values for the features with the highest ground truth standard deviation across n = 8 non-pregnant controls (indicated by arrows in d).

To validate SIMCO’s digital twin models, we compared simulated immune responses with the observed, real-world responses in the perturbation atlas. Models were evaluated based on three metrics using a 4-fold cross-validation strategy: the mean absolute error (MAE) and the coefficient of determination (R^2^) between simulated and observed population-level medians as well as the maximum mean discrepancy (MMD) between simulated and observed marker distributions. Example evaluations are shown for the responses to IL-2/4/6 stimulation (**Figure 4b**). Performance summary metrics for all stimulation conditions are provided in **Supplementary Table S2**. For benchmarking, we included the identity baseline (simply returning untreated distributions) as well as the observed training baseline (simply returning the perturbed distributions of the training patients). Across all metrics, SIMCO consistently outperformed the training baseline, validating that it captures patient heterogeneity and enables accurate individualized perturbation modeling (**Figure 4b**).

Comparing simulated and observed marker distributions for example features with strong IL-2/4/6-induced stimulation response (> 0.1 arcsinh difference) highlighted the accuracy of SIMCO’s digital twin models for heterogeneous single-cell behavior (**Figure 4c**). The MAE of single-cell simulations generally increased with stronger signaling response (**Figure 4d**). The simulation error correlated with interpatient variability in the observed signaling responses (Spearman’s R = 0.93, **Figure 4d**). However, SIMCO simulations still contained relative differences between patients even for the most variable responses (**Figure 4e**).

### SIMCO predicts treatment effects on time to labor

The main purpose of the SIMCO framework is to leverage single-cell-level digital twins to predict how immunomodulatory treatments influence clinical outcomes. Using an independent mass cytometry dataset collected by Stelzer et al. 2021 over the last 120 days of pregnancy, SIMCO generated digital twins for 177 blood samples collected in term (153 samples, 58 patients) and preterm (14 samples, 5 patients) pregnancies (**Figure 5a**).^33^ We first established that a sparse multivariable model built on the untreated single-cell-digital twin dataset accurately predicted the clinical outcome time to labor (Pearson R = 0.64, root mean squared error [RMSE] = 24.1 days, term: 24.0 days, preterm 25.4 days, **Figure 5b**). Furthermore, after estimating the expected pregnancy duration by combining the time to labor prediction with the sampling timepoint, SIMCO accurately differentiated term from preterm digital twins (AUROC = 0.73 [95% confidence interval = 0.6–0.85], **Figure 5c, Supplementary Table S3**).

**Figure 5:**
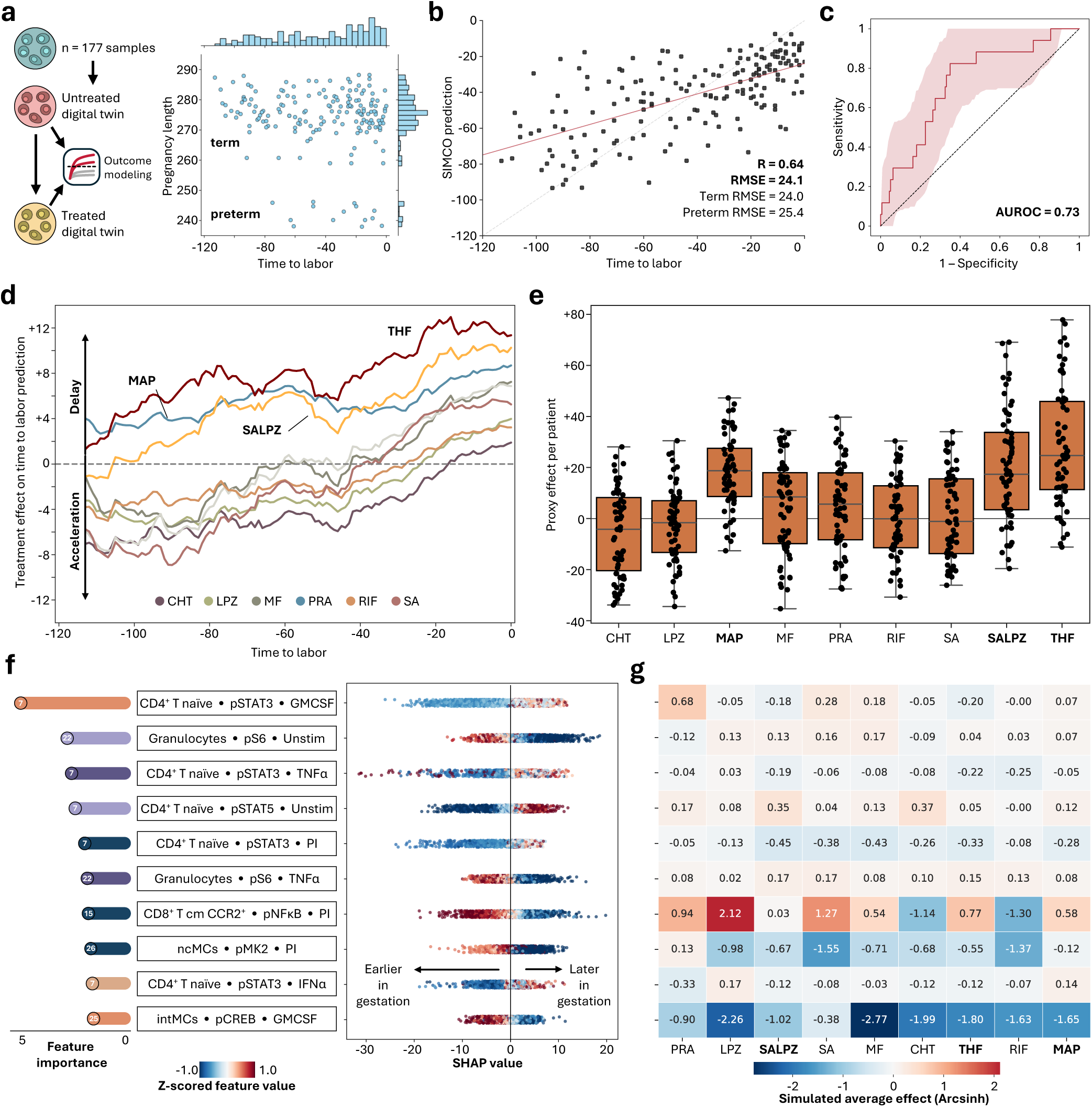
SIMCO predicts treatment effects on time to labor. **a.** SIMCO generated treated and untreated digital twins for the independent study from Stelzer et al. The study cohort included 177 samples from 63 patients collected during the second and third trimesters of pregnancy (including 14 samples from 5 patients with sPTL). The primary outcome was the time to spontaneous labor onset. **b.** SIMCO’s performance to predict time to labor. Pearson R and root mean squared error (RMSE) are shown. **c.** Area under the receiver operating characteristic curve (AUROC) plot for identifying preterm samples via the expected pregnancy length, calculated from SIMCO’s time to labor prediction and the sampling timepoint. **d.** Time-dependency (rolling average of 30 days) of SIMCO-predicted treatment effects across the last 120 days of pregnancy. **e.** Proxy effect per patient (n = 63) on the gestational length for nine candidate drugs. **f.** Top 10 model features (out of 21 [18–30] features total) in descending order of mean Shapley Additive exPlanations (SHAP) value as a measure of feature importance in the SIMCO simulation. The SHAP summary plot reveals temporal association of z-scored feature values with gestational age. **g.** Heatmap of average drug effects (n = 177) on the top 10 model features, as simulated by SIMCO. Candidate drugs are abbreviated as follows: chlorthalidone (CHT), maprotiline (MAP), metformin (MF), lansoprazole (LPZ), pravastatin (PRA), rifabutin (RIF), salicylic acid (SA), salicylic acid + lansoprazole (SALPZ), and 5-methyltetrahydrofolate (THF).

SIMCO modeled the effect of the candidate drugs on the clinical outcome “time to labor” (see **Methods**): For each simulated sample, SIMCO calculated the difference in the predicted time to labor between candidate drug-treated and untreated digital twins (**Figure 5d**). Maprotiline, THF, and SALPZ consistently delayed the predicted time to labor throughout all measured gestational time points. In contrast, other candidate drugs displayed a biphasic effect, first accelerating then decelerating the progression of gestation. To capture the cumulative effect of treatment over time, we aggregated data from the same patient across all three sampling time points (**Figure 5e**). The strongest proxy effect on gestational length was achieved by THF (24.7 days, interquartile range [IQR] = 11.4–45.9), maprotiline (18.8 days, IQR = 8.6–27.5), and SALPZ (17.3 days, IQR = 3.5–33.8). Reflecting clinical reality, not all patients were modeled to benefit from a given treatment: THF, SALPZ, and maprotiline were predicted to prolong gestational length in 90.5%, 87.3%, and 82.5% of patients. Other candidate drugs, such as SA alone, only profited 46% of the patients.

The sparse prediction model for time to labor used 21 key informative immune features. Of the top 10 model features, five represented CD4^+^ T cell signaling responses: notably, reduced pSTAT3 levels in response to GMCSF, TNFα, PI, and IFNα was associated with earlier gestational time points, i.e., STAT3 activation increased with approaching labor (**Figure 5f**). In the innate immune compartment, pS6 responses in granulocytes showed dynamic modulation over time, with both baseline and TNFα-induced pS6 levels decreasing as pregnancies approached labor.

SIMCO also enables mechanistic insight into individual drug effects: To delay the predicted onset to labor, candidate drugs shifted key immune features of the predictive model toward patterns associated with earlier pregnancy stages. THF and SALPZ did so by consistently suppressing pSTAT3 signaling in CD4^+^ naïve T cells and enhancing pS6 responses in granulocytes, thereby counteracting immunological signatures linked to pregnancy progression (**Figure 5g**). Maprotiline and THF also boosted pNFκB signaling after PI stimulation in CD8^+^ CCR2^+^ central memory T cells, which was higher in earlier gestation. However, some drug effects paradoxically shifted features towards patterns associated with later pregnancy stages. For instance, THF and SALPZ, strongly inhibited pMK2 activation in non-classical monocytes (ncMCs) following PI stimulation and pCREB activation in intermediate monocytes (intMCs) following GMCSF stimulation, whereas decreased activation in these features would normally be expected closer to labor onset. Similarly, both chlorthalidone and SALPZ induced baseline activation of pSTAT5 in CD4^+^ naïve T cells, a feature the model associated with later stages of pregnancy. However, SIMCO intrinsically integrates and prioritizes these multiple effects when generating outcome predictions for the treated digital twin. By integrating many immunomodulatory effects across cell types, stimuli, and signaling pathways, SIMCO provides a quantitative, systems-level evaluation of each candidate drug’s net impact on gestational timing.

## DISCUSSION

SIMCO is a single-cell-level digital twin framework developed to model clinical outcome effects of immunomodulatory candidate drugs. In a first step, SIMCO learns immune mechanisms based on a high-dimensional single-cell perturbation dataset. In a second step, the trained model is parametrized with single-cell data of unseen individuals. By contrasting treated and untreated digital twin models, SIMCO quantifies how interventions alter clinical outcomes while offering unique single-cell-level mechanistic explainability. We showcase the application of SIMCO for sPTL prevention, a setting where in silico clinical trials are especially valuable given the constraints of interventional studies in pregnancy. Simulating treatment with nine preselected candidate drugs for the prevention of sPTL, SIMCO identified THF, maprotiline, and SALPZ as strong candidates to delay labor onset in preterm pregnancies.

We employed SIMCO with the premise that altering labor-associated immune signatures could delay labor onset and therefore prevent sPTL. By quantitatively evaluating drug-induced shifts of simulated immune states, SIMCO identified candidate drugs that stabilize, or reverse immunological changes associated with the gestational progression towards labor. Of note, the effects were quantified based on a single-dose exposure; in practice, repeated administration (e.g., daily) would likely produce accumulating effects. Aggregation of the drug effects on the patient level (across up to three samples) was therefore used as a proxy to continuous treatment. The three most effective drugs or drug combinations in delaying predicted time to labor were THF, maprotiline, and SALPZ. Of these, SALPZ, the combination of low-dose aspirin (81 mg) and lansoprazole (30 mg), is of particular interest: Aspirin treatment alone, when initiated between six and 12 weeks of gestation, has demonstrated modest reductions in the incidence of sPTL in large clinical trials.^25,46,47^ However, the small effect size has prompted ongoing debate around optimal dosing, timing, and the potential for improved efficacy through combination with other agents. Lansoprazole, a proton pump inhibitor with known anti-inflammatory properties, inhibits LPS-induced, TLR4-mediated NFκB activation,^48^ a pathway implicated in inflammation-driven preterm labor.^49^ In a murine model of LPS-induced fetal wastage, lansoprazole treatment significantly improved fetal survival.^23^ Our findings further suggest that SA and lansoprazole act synergistically, particularly in the modulation of JAK-STAT signaling. This synergy may enhance their immunomodulatory effect, providing a rationale for clinical evaluation of the combination for their potential to reduce sPTL incidence.

THF and maprotiline also delayed the simulated time to labor. While current guidelines recommend folic acid supplementation primarily for the prevention of neural tube defects,^50^ results from large retrospective studies show that folic acid supplementation is associated with a reduced risk of preterm delivery.^51^ Our digital twins suggest that folic acid/THF supplementation may benefit individuals at high risk for sPTL by delaying their immunological progression toward labor. Maprotiline is a discontinued tetracyclic antidepressant used for treatment of major depressive disorder and anxiety. Per FDA guidelines, maprotiline was a category B drug that could be continued throughout gestation with no evidence of increased risk of sPTL.^52,53^ However, maprotiline use often comes with significant side effects such as fatigue, weight gain, and anticholinergic effects. In addition, maprotiline also seems to mediate diverse immunomodulatory effects; however, its appeal as a preventive strategy for reducing sPTL risk is low due to its side effect profile and the general controversy around the use of antidepressants during pregnancy.

A key strength of the SIMCO framework is the ability to monitor simulated drug effects across the entire immune system and comprehensively weigh desirable and undesirable effects on the clinical outcome. Drugs that delayed the predicted time to labor predominantly modulated signaling in CD4^+^ T cells and granulocytes, both of which are tightly regulated throughout gestational progression. Notably, suppression of pSTAT3 responsiveness in naïve CD4^+^ T cells can promote immune tolerance and limit proinflammatory T cell differentiation.^54,55^ A sustained reduction in pSTAT3 activation might therefore protect against sPTL.^56–58^ Similarly, enhanced pSTAT5 responsiveness, which is central to the development of regulatory T cells,^59,60^ may support maternal-fetal tolerance. These subtle shifts in CD4^+^ naïve T cell homeostasis observed in samples treated with THF, SALPZ, or maprotiline are highly relevant to maintaining appropriate gestational immune adaptations.

Innate immune cells also show immunosuppressive adaptations to sustain pregnancy, before taking part in the proinflammatory initiation of labor.^61–63^ In this context, drugs that bolstered pS6 activity in granulocytes, e.g., metformin, SA, and SALPZ, beneficially influenced time to labor predictions in our simulations. While mTOR signaling is complex, activation of mTOR/pS6 activity has been shown to enhance IL-10 production and anti-inflammatory (M2-like) polarization in myeloid immune cells.^64,65^ Drugs that maintain higher mTOR pathway activity mid-pregnancy may thus help preserve immune tolerance and suppress labor-associated inflammation.

SIMCO enables meaningful data augmentation by extrapolating from unstimulated samples to infer responses under unmeasured conditions, thereby generating digital twins of the immunome. This approach assumes that the perturbation atlas is informative for the clinical dataset, as mismatched datasets from different patient populations could invalidate such extrapolations.^35^ While OT methods have seen growing use in basic science applications such as trajectory inference^66–68^ and perturbation modeling,^69–71^ their adoption in translational research remains limited. Here, we showcase the utility of OT-based modeling within SIMCO to simulate gestational immune states and model outcome-level treatment effects. The strong predictive performance of the generated digital twins, approaching that of the original mass cytometry dataset from Stelzer et al., indicates that SIMCO accurately models immune mechanisms underlying labor timing and bases its prediction of treatment effects on relevant immune responses.

This study has several limitations. Although our immune perturbation atlas was profiled at the single-cell level, it included a relatively small number of individuals during the second trimester and focused exclusively on peripheral blood, rather than tissue-resident immune populations. This design reflects our goal of developing a minimally invasive, blood-based assay that can be scaled and incorporated in biomarker-driven clinical trial designs; however, it may overlook key tissue-specific immune dynamics involved in the pathophysiology of sPTL. Additionally, our in vitro experiments required selecting specific compounds at specific concentrations for testing. Several of the investigated drugs are metabolized in the liver to active and inactive metabolites, a process that cannot be replicated in vitro. To approximate physiological relevance, we selected the metabolites with the highest plasma concentrations following oral administration; for instance, for aspirin and folic acid, we tested SA and THF, respectively. While we cannot exclude the possibility that other metabolites also exert immunomodulatory effects, our experimental design prioritized those most likely to reflect systemic exposure. Effective concentrations strictly followed observed plasma levels after oral administration of clinically relevant doses, except for THF, where we chose a high-dose (4 mg), as many patients reported taking prenatal vitamins (including THF at the recommended daily value). Moreover, while our simulation approach allows for prioritizing which treatment to test further, in vivo validation in appropriate animal models or human clinical studies will be essential to confirm therapeutic efficacy. Finally, it remains unclear whether the peripheral immune changes we observe are causal drivers or downstream consequences of sPTL pathophysiology. Our work assumes that peripheral immune features associated with time to labor influence pregnancy progression. Yet, it is important to recognize that candidate drugs exert additional effects beyond the model features, which could also contribute to or oppose therapeutic benefit.

In conclusion, we present SIMCO as a proof-of-concept framework for single-cell-level digital twins that simulate immune responses and model outcome-level treatment effects. With the ability to generate digital twins from baseline immune profiles of unseen individuals, SIMCO offers a scalable and minimally invasive addition to traditional clinical trials for initial compound prioritization. Our work represents a critical step towards data-driven, immune-guided therapeutic strategies for sPTL prevention. Future directions should include real-time immune monitoring, expansion of perturbation libraries, and clinical validation of treatment effects. Ultimately, SIMCO offers a new path toward precision obstetrics by aligning immunological insight with computational modeling and personalized care.

## METHODS

### Study participant details and experimental setup

#### Study design

The aim of this in vitro drug screen was to identify reproducible immunomodulatory effects of select candidate drugs for the prevention of sPTL. The study was conducted at Stanford Hospital (Stanford, CA, USA) and approved by the Institutional Review Board (protocol number 68417). All participants provided informed consent. Non-pregnant volunteers were recruited during the first week of their menstrual cycle. Inclusion criteria included age 18 – 45 years, BMI < 40, and no ongoing immunomodulatory treatment. Pregnant individuals were recruited between 20 and 27 weeks of gestation if they were 18 – 45 years old, had a pre-pregnancy BMI < 40, were carrying a singleton pregnancy, and without prior pregnancies (G1P0). Exclusion criteria included fertility treatment, chronic conditions outside of pregnancy (e.g. hypertension, diabetes), concurrent immunomodulatory medication, diagnosed pregnancy disorders and fetal anomalies, as well as autoimmune diseases (e.g. lupus, rheumatoid arthritis, Sjogren’s, or Hashimoto’s). A single blood draw of 12 mL was collected in heparinized tubes. In total, eight non-pregnant controls and eleven pregnant individuals participated in the study. Researchers conducting the analyses were not blinded. Basic demographics and pregnancy characteristics are summarized in **Table 1**.

#### Ex vivo drug screening assay

Whole blood was collected from study subjects in heparinized tubes and processed between 30 and 60 min after blood draw. The drug screen assessed 96 conditions per sample and was automated using an Opentrons Flex pipetting robot. Drug aliquots were prepared as 200x stocks from powder dissolved in DMSO (except 5-THF, 100x stock in 1:1 water:DMSO). In a 96-well deep-well block, RPMI with 2x drug concentrations was added and warmed to 37 °C. Heparinized whole blood was added 1:1 to the RPMI/drug dilution in each well, achieving the final 1x concentration for each drug (0.5 % DMSO). The final drug concentrations per drug can be found in **Table 2**. The pretreated whole blood was incubated for 1 h at 37 °C and 5 % CO_2_ in a humidified incubator on an orbital shaker. After 1h, the samples were stimulated with LPS (50 ng/ml; InvivoGen, San Diego, CA), IFNα (50 ng/ml; PBL Assay Science, Piscataway, NJ), IL-33 (50 ng/ml; R&D Systems, Minneapolis, MN), GM-CSF (50 ng/ml; R&D Systems), TNFα (50 ng/ml; R&D Systems), a cocktail of IL-2, IL-4, and IL-6 (each 50 ng/ml; R&D Systems), PI (0.5×, ThermoFisher, Waltham, MA), or left unstimulated. After a 2 h incubation at 37 °C and 5 % CO_2_ in a humidified incubator on an orbital shaker, the whole blood culture was fixed with proteomic stabilizer (SMART TUBE Inc., San Carlos, CA) for 20 min. Samples were treated with blood cell lysis buffer twice for 5 min, then washed and stored at -80 °C until further processing.

#### 96-plex layout and barcoding

All 96 screening conditions for each donor were organized in a standardized 12 × 8 format in a 2 ml deep-well block. Stimulation conditions were organized as rows, treatment conditions as columns. Three DMSO controls were included in column one, six, and 12, the candidate drugs were organized as depicted in **Figure 2a**. A 96-plex doublet-removing barcoding scheme was optimized to allow pooling of all conditions together for mass cytometry staining and acquisition to avoid batch effects.^72^ Mass-tag barcodes were designed using the isotopes Pd104, Pd105, Pd106, Pd108, Pd110, Pt194, Pt195, Pt196, and Au197. Isotope concentrations were titrated to 10 nM (Au, Pt isotopes) and 100 nM (Pd isotopes) to achieve optimal barcode separation. Out of the 126 barcodes available in a choose-4-out-of-9 scheme, the 96 barcodes with the highest purity (based on the purity of the underlying isotopes) were selected, while the remaining 30 were used to monitor the rate of wrongly assigned cells (**Supplementary Figure S3**). Barcode reagents were prepared in DMSO and used after transient permeabilization with 0.02 % saponin/PBS.^73^ Across all 19 blood donors, the average barcode assignment rate with a threshold of 0.1 was 71.8 % (± 4.2 %). For each condition, 100 µl of blood yielded on average 138.0 × 10^3^ (118.2 -162.1 × 10^3^) live leukocytes for pregnant donors and 94.6 × 10^3^ (76.9 -111.4 × 10^3^) for non-pregnant donors.

#### Opentrons Flex operation

Protocol operation with the Opentrons Flex require an eight-channel and a single-channel pipette. The protocol for the in vitro drug exposure and whole blood lysis required around 5 h of operation. A custom tube rack and a pipette basin holder were 3D printed to allow for the used deck configuration. The protocol for 96 well barcoding required around 2h of operation. The protocols as well as the .stl files for the custom parts are available on the GitHub repository.

#### Antibody staining and mass cytometry analysis

Pooled barcoded samples from all 96 conditions (40 × 10^6^ cells) were incubated with Fc-block (Human TruStain FcX, Biolegend) for 10 min, then stained with a master mix of surface antibodies for 30 min in cell staining medium (PBS with 0.5 % BSA and 0.02 % sodium azide). After washing, the cells were permeabilized in ice-cold 100 % methanol, washed, and stained for 30 min with a master mix of intracellular antibodies (**Supplementary Table S1**). Following intracellular staining, cells were washed and resuspended in an iridium intercalator (Standard Biotools) solution containing 2 % paraformaldehyde. Finally, samples were washed with ddH2O and pelleted. The data was acquired using a mass cytometry XT instrument (Standard Biotools). The resulting mass cytometry data were normalized across each run using 6-EQ normalization beads. Debarcoding of the data was achieved using the Single Cell Debarcoder v0.2 in Matlab at a barcode separation threshold of 0.1.^72^

#### Stimulation responses and drug-induced effects

Using CellEngine (Primity Bio), cell populations were defined with manual gating (see **Supplementary Figure S2**). Expression levels of intracellular phospho-signaling markers (pSTAT1, pSTAT3, pSTAT4, pSTAT5, pSTAT6, pMAPKAPK2, pCREB, pPLCγ2, pS6, pERK1/2, pP38, pNFκB, and total IκB) and cell surface functional markers (PD-1, PD-L1, CD25, CD36, CD44, GLUT1, HLA-DR) were exported as population median per stimulation condition, drug treatment, and study participant. Per individual, the combination of eight stimulation × 28 immune cell subsets × 20 functional proteins produced a total of 4480 features for each of the nine drug treatments and the three DMSO controls. The extracted dataset was arcsinh transformed with a cofactor of 5. The DMSO control triplicate was used to determine the intra-assay variability. Per patient per feature, a smooth spline was fitted across all columns (k = 2, smoothing = 1.5, weight = 1 for drug-columns, weight = 2 for DMSO columns) to account for time-dependent effects in signaling response (see **Supplementary figure S6**). Drug effects were calculated as difference to the spline-modeled column-specific corrected DMSO control within the same stimulation condition. Feature stimulation effects were classified as strong response if the arcsinh difference to unstimulated was > 0.1 (0.01 for IL-33). For 168 out of 196 cell type/stimulation combination, at least one strong stimulation response was observed.

### Simulated Immunome Modeling of Clinical Outcomes (SIMCO) workflow

#### Overview

We developed SIMCO to rapidly evaluate treatment effects of immunomodulatory drugs on a clinical outcome via single-cell simulations. The prediction ensemble utilizes CellOT to simulate single-cell immune states and Stabl for feature selection and outcome modeling. A dataset of high-dimensional perturbation experiments (e.g., from mass cytometry) should be generated in a relevant patient population, evaluating immune cell responses when challenged with s stimulatory ligands in the presence or absence of t candidate drugs (**Supplementary Figure S1a**). After identifying cell types (e.g., using manual gating), cell type-specific CellOT models are trained for each stimulation scenarios s; separately, for each stimulation condition, CellOT models for each drug treatment scenarios t (**Supplementary Figure S1b**). The learned single-cell behavior is then projected onto a separate immunome dataset with documented outcomes and matching single-cell readouts (e.g., antibody panel). For n unstimulated samples with p unstimulated features, responses are simulated to s stimulation scenarios, creating a feature matrix of p × (s + 1) features. This expanded feature matrix is then also transformed into t “treated” feature spaces (**Supplementary Figure S1c**). Outcome modeling using the Stabl algorithm is performed on the untreated, simulation-expanded data set in cross-validation: Per cross-validation fold, a predictive model is built on the training cohort to then predict the outcome on a hold-out test set of patients as well as their “treated” twins (**Supplementary Figure S1d**). The global model performance is evaluated across folds; treatment effects were evaluated per patient per cross-validation fold as the difference between the predictions on untreated in treated (**Supplementary Figure S1e**). The final model built on the entire dataset can be used to identify patients at risk in an unseen cohort. By using a specific model prediction as treatment target (continuous or categorical), the model can also steer personalized treatment strategies (**Supplementary Figure S1f**).

#### Single-cell simulation using CellOT

CellOT was introduced by Bunne et al. 2023 as an implementation of OT learning using an input convex neural network with four fully connected hidden layers. We adapted the network architecture to include a four-layer bottleneck structure (128, 128, 64, 32). We reduced the latent-space dimensions from 50 to 42 to prevent overfitting and stopped training at 50,000 epochs (instead of 100,000). CellOT models were trained on single-cell data from individual cell types, identified by manual gating, as. CellOT models trained per cell type achieved better predictive accuracy than CellOT models built on all leukocytes together.

#### Single-cell simulation performance evaluation

We evaluated CellOT’s single-cell simulation performance using four-fold cross-validation. For performance evaluation, CellOT models were trained in four-fold cross-validation. For each fold, CellOT was trained on a subset of patients, and predictions were generated for the held-out patients. Predicted single-cell data were summarized by calculating population medians per marker. To contextualize the performance, we compared CellOT against two simple baselines. In the identity baseline, predictions were set equal to the untreated (baseline) distribution of the same patient, representing the trivial case of assuming no perturbation effect was learned by CellOT. In the training baseline, predictions were set equal to the observed perturbed distributions from the training patients, representing the case of directly reusing available training data without modeling new responses. Performance metrics were then computed consistently across models: mean absolute error (MAE) between predicted and observed population medians; maximum mean discrepancy (MMD) between predicted and observed marker distributions per cell type, marker, and patient; and R² values quantifying the correlation between predicted and observed population medians across markers for each cell type and patient.

#### Final single-cell simulation model

For the final analysis, CellOT models were trained using all 19 patients (pregnant and non-pregnant) from the perturbation dataset. For stimulation response models, we trained a separate CellOT model for each of the 168 cell type-stimulation combinations that showed at least one strong stimulation response. For treatment effect models, we extended the training to predict drug effects on the same 168 cell type-stimulation combinations, as well as drug effects on all 28 cell types at unstimulated baseline.

#### Independent dataset with time to labor outcome

We used a longitudinal mass cytometry dataset recently published by our group as clinical reference dataset for simulation experiments.^33^ The dataset comprised 177 samples from 63 participants enrolled during the last 120 days of an uncomplicated, singleton pregnancy. Between one and three blood samples were collected throughout this period until patients went into spontaneous labor. For simulation experiments, only the 177 unstimulated samples (15 min, 37 °C) were used (original stimulation conditions included unstimulated, IL-2/4/6, LPS, IFNα, and GMCSF). The antibody panel largely overlapped, and manual gating was adjusted to match the perturbation dataset completely. Population medians were exported and arcsinh transformed with a cofactor of 5.

#### Multivariable outcome modeling using Stabl

The Stabl algorithm allows for the selection of reliable biomarkers from multiomic datasets.^32^ As input data, population medians of baseline expression levels (unstimulated) or stimulation responses (arcsinh difference between stimulated and unstimulated) were used. To account for the longitudinal nature of the dataset, we used patient shuffle split to assess the robustness of our model. We applied a penalization matrix to remove weak stimulation responses (median response across all samples < 0.1 arcsinh difference) from the input feature set. This penalization resulted in a dataset size of 364 unstimulated features (28 cell types × 13 markers) and 691 stimulation response features (out of 7 stimulation conditions × 28 × 13 = 2,548 features). Using a grid search across multiple hyperparameters (**Supplementary Table S4**, all other hyperparameters used as default), we selected the Stabl parameters providing the best feature set based on the RMSE. For the final model fit, we then iterated over a grid search for XGBoost hyperparameters optimizing the AUROC (**Supplementary Table S4**).

#### Evaluation of simulated drug effects

Within the cross-validation strategy used to assess model performance, we calculated the time to labor predictions for both the held-out test patients as well as their “treated” counterparts. To reduce the effect of modeling error on the treatment evaluation, we compared the model prediction on the “treated” patient data to the same patient’s “no drug” model prediction.

#### Computational resources

CellOT was executed using the Python package ‘CellOT’ (version 0.1). CellOT training was performed on the Stanford-provided cloud computing platform Sherlock with maximum RAM request of 32 GB, maximum job time of 20 h, maximum 16 CPU per job. Stabl was run using the Python package ‘Stabl lw’. Basic data analysis steps, including time-dependent variability correction and dataset penalization, were executed using the Python packages ‘SciPy’ (version 1.10.4), ‘scikit-learn’ (version 1.5.2), ‘pandas’ (version 2.2.3) and ‘numpy’ (version 1.26.4). For visualization, we used ‘seaborn’ (version 0.13.2) and ‘matplotlib’ (version 3.10.3).

#### Univariate analysis

Statistical comparisons between non-pregnant and pregnant individuals were calculated using a Mann-Whitney U test with false discovery rate (FDR) correction (FDR < 0.05, Benjamini-Hochberg). Comparisons between treatment and DMSO-control were calculated as paired student’s t test with Benjamini-Hochberg correction (FDR < 0.05). Pregnancy specificity of drug effects was calculated as the absolute difference in effect size between pregnant and non-pregnant donors, normalized by the total absolute effect magnitude across both groups.

## Supplementary Materials

Figure S1: Detailed SIMCO workflow.

Figure S2: Gating strategy.

Figure S3: 96 well barcoding.

Figure S4: All drug effect heatmaps.

Figure S5: Comparison of SA, LPZ, and SALPZ drug effects.

Figure S6: DMSO modeling.

Table S1: Antibody panel.

Table S2: CellOT performance in cross-validation across stimulation conditions.

Table S3: Predictive performances for SIMCO-simulated data, Stelzer et al., and EGA.

Table S4: Grid search parameters for Stabl/XGBoost models.

## Acknowledgments

We thank all individuals for their participation in this study. We thank Dr. Yasser El Sayed for his thoughtful input and guidance on the analyses. Parts of figures 1 and 2 were created using biorender.com.

## Author contributions

Conceptualization, BG, JE, MSato, IAS, LG, and DKS. Clinical data collection, KA, AA, and KM. Formal analysis, JE, PN, OF, JNA, VB, and MSabayev. Data acquisition, JE, EAG, DF, EOK, MSato, AST, MD, AC, and DKG. Methodology, JE, JH, TTO, MS, LG, NA, GMS, YJB, RJW, DJL, DKS, and BG. Supervision, MS, LG, DKS, and BG. Visualization, JE, PN, OF, JNA, RL, EK, and VB. Writing – original draft, JE, PN, MSato, OF, DKG, and BG. Writing – review & editing, BG, JE, and all authors.

## Lead contact

Further information and requests for resources and reagents should be directed to and will be fulfilled by the lead contact, Brice Gaudillière.

## Materials availability

All materials used in this study are commercially available, as specified in the supplementary tables. This study did not generate new unique reagents.

## Data and code availability

The preprocessed data of the perturbation dataset can be found as pre-gated FCS files on dryad.com: https://doi.org/10.5061/dryad.sqv9s4ngm. The source code required to reproduce analysis steps is publicly available on Github: https://github.com/ofondeur/SIMCO/. Any additional information required to reanalyze the data reported in this paper is available from the lead contact upon request.

## Clinical trial number

not applicable

## References

1. Liu, L. et al. Global, regional, and national causes of under-5 mortality in 2000–15: an updated systematic analysis with implications for the Sustainable Development Goals. The Lancet 388, 3027–3035 (2016).

2. Goldenberg, R. L., Culhane, J. F., Iams, J. D. & Romero, R. Epidemiology and causes of preterm birth. Lancet 371, 75–84 (2008).

3. Romero, R., Dey, S. K. & Fisher, S. J. Preterm labor: One syndrome, many causes. Science 345, 760–765 (2014).

4. Han, X. et al. Differential Dynamics of the Maternal Immune System in Healthy Pregnancy and Preeclampsia. Front. Immunol. 10, 1305 (2019).

5. Aghaeepour, N., et al. An immune clock of human pregnancy. Sci. Immunol. 2, eaan2946 (2017).

6. Tarca, A. L. et al. The prediction of early preeclampsia: Results from a longitudinal proteomics study. PLoS ONE 14, e0217273 (2019).

7. Gomez-Lopez, N. et al. The Cellular Transcriptome in the Maternal Circulation During Normal Pregnancy: A Longitudinal Study. Front. Immunol. 10, 2863 (2019).

8. Ngo, T. T. M. et al. Noninvasive blood tests for fetal development predict gestational age and preterm delivery. Science 360, 1133–1136 (2018).

9. Racicot, K., Kwon, J., Aldo, P., Silasi, M. & Mor, G. Understanding the Complexity of the Immune System during Pregnancy. American J Rep Immunol 72, 107–116 (2014).

10. Pique-Regi, R. et al. Single cell transcriptional signatures of the human placenta in term and preterm parturition. eLife 8, e52004 (2019).

11. Tarca, A. L. et al. Crowdsourcing assessment of maternal blood multi-omics for predicting gestational age and preterm birth. Cell Reports Medicine 2, 100323 (2021).

12. Rowe, J. H., Ertelt, J. M., Xin, L. & Way, S. S. Pregnancy imprints regulatory memory that sustains anergy to fetal antigen. Nature 490, 102–106 (2012).

13. Erlebacher, A. Mechanisms of T cell tolerance towards the allogeneic fetus. Nat Rev Immunol 13, 23–33 (2013).

14. Tsuda, S., Nakashima, A., Shima, T. & Saito, S. New Paradigm in the Role of Regulatory T Cells During Pregnancy. Front. Immunol. 10, 573 (2019).

15. Somerset, D. A., Zheng, Y., Kilby, M. D., Sansom, D. M. & Drayson, M. T. Normal human pregnancy is associated with an elevation in the immune suppressive CD25+ CD4+ regulatory T-cell subset. Immunology 112, 38–43 (2004).

16. Gomez-Lopez, N., Tanaka, S., Zaeem, Z., Metz, G. A. & Olson, D. M. Maternal circulating leukocytes display early chemotactic responsiveness during late gestation. BMC Pregnancy Childbirth 13, S8 (2013).

17. Farias-Jofre, M. et al. Pregnancy tailors endotoxin-induced monocyte and neutrophil responses in the maternal circulation. Inflamm Res 71, 653–668 (2022).

18. Mor, G., Aldo, P. & Alvero, A. B. The unique immunological and microbial aspects of pregnancy. Nat Rev Immunol 17, 469–482 (2017).

19. Gomez-Lopez, N. et al. Inflammasomes: Their Role in Normal and Complicated Pregnancies. The Journal of Immunology 203, 2757–2769 (2019).

20. Gomez-Lopez, N. et al. The immunobiology of preterm labor and birth: intra-amniotic inflammation or breakdown of maternal–fetal homeostasis. Reproduction 164, R11–R45 (2022).

21. Miller, D. et al. Single-Cell Immunobiology of the Maternal-Fetal Interface. J Immunol 209, 1450–1464 (2022).

22. Feyaerts, D. et al. The single-cell immune profile throughout gestation and its potential value for identifying women at risk for spontaneous preterm birth. Eur J Obstet Gynecol Reprod Biol X 25, 100371 (2025).

23. Le, B. L., Iwatani, S., Wong, R. J., Stevenson, D. K. & Sirota, M. Computational discovery of therapeutic candidates for preventing preterm birth. JCI Insight 5, (2020).

24. Feyaerts, D. et al. Predicting Spontaneous Preterm Birth Using the Immunome. Clin Perinatol 51, 441–459 (2024).

25. Hoffman, M. K. et al. Low-dose aspirin for the prevention of preterm delivery in nulliparous women with a singleton pregnancy (ASPIRIN): a randomised, double-blind, placebo-controlled trial. Lancet 395, 285–293 (2020).

26. Prediction and Prevention of Spontaneous Preterm Birth: ACOG Practice Bulletin, Number 234. Obstetrics & Gynecology 138, e65–e90 (2021).

27. Society for Maternal-Fetal Medicine Statement: Response to the Food and Drug Administration’s withdrawal of 17-alpha hydroxyprogesterone caproate. American Journal of Obstetrics and Gynecology 229, B2–B6 (2023).

28. Manuck, T. A., Gyamfi-Bannerman, C. & Saade, G. What now? A critical evaluation of over 20 years of clinical and research experience with 17-alpha hydroxyprogesterone caproate for recurrent preterm birth prevention. Am J Obstet Gynecol MFM 5, 101108 (2023).

29. Pappalardo, F., Russo, G., Tshinanu, F. M. & Viceconti, M. In silico clinical trials: concepts and early adoptions. Brief Bioinform 20, 1699–1708 (2019).

30. Laubenbacher, R. et al. Building digital twins of the human immune system: toward a roadmap. *npj Digit*. Med. 5, 64 (2022).

31. Bendall, S. C. et al. Single-cell mass cytometry of differential immune and drug responses across a human hematopoietic continuum. Science 332, 687–696 (2011).

32. Hédou, J. et al. Discovery of sparse, reliable omic biomarkers with Stabl. Nat Biotechnol (2024) doi:10.1038/s41587-023-02033-x.

33. Stelzer, I. A. et al. Integrated trajectories of the maternal metabolome, proteome, and immunome predict labor onset. Sci Transl Med 13, eabd9898 (2021).

34. Bodenmiller, B. et al. Multiplexed mass cytometry profiling of cellular states perturbed by small-molecule regulators. Nat Biotechnol 30, 858–867 (2012).

35. Bunne, C., Schiebinger, G., Krause, A., Regev, A. & Cuturi, M. Optimal transport for single-cell and spatial omics. Nat Rev Methods Primers 4, 58 (2024).

36. Bunne, C. et al. Learning single-cell perturbation responses using neural optimal transport. Nat Methods 20, 1759–1768 (2023).

37. Shah, J. V., Patel, D. P., Shah, P. A., Sanyal, M. & Shrivastav, P. S. Simultaneous quantification of atenolol and chlorthalidone in human plasma by ultra-performance liquid chromatography–tandem mass spectrometry. Biomedical Chromatography 30, 208–216 (2016).

38. Maguire, K. P., Norman, T. R., Burrows, G. D. & Scoggins, B. A. An evaluation of maprotiline intravenous kinetics and comparison of two oral doses. Eur J Clin Pharmacol 18, 249–254 (1980).

39. Eyal, S. et al. Pharmacokinetics of Metformin during Pregnancy. Drug Metab Dispos 38, 833–840 (2010).

40. Song, M., Gao, X., Hang, T.-J. & Wen, A.-D. Pharmacokinetic properties of lansoprazole (30-mg enteric-coated capsules) and its metabolites: A single-dose, open-label study in healthy Chinese male subjects. Current Therapeutic Research 70, 228–239 (2009).

41. Costantine, M. M. et al. Safety and pharmacokinetics of pravastatin used for the prevention of preeclampsia in high-risk pregnant women: a pilot randomized controlled trial. American Journal of Obstetrics & Gynecology 214, 720.e1–720.e17 (2016).

42. Narang, P. K., Lewis, R. C. & Bianchine, J. R. Rifabutin absorption in humans: Relative bioavailability and food effect. Clinical Pharmacology & Therapeutics 52, 335–341 (1992).

43. Rc, D. et al. Comparative Bioavailability Study of Two 81 mg Coated Tablet Formulations of Acetylsalicylic Acid in Fasting Healthy Volunteers. J Bioequiv Availab 09, (2017).

44. Zheng, X.-H., Jiang, L.-Y., Zhao, L.-T., Zhang, Q.-Y. & Ding, L. Simultaneous quantitation of folic acid and 5-methyltetrahydrofolic acid in human plasma by HPLC–MS/MS and its application to a pharmacokinetic study. J Pharm Anal 5, 269–275 (2015).

45. Abu-Raya, B., Michalski, C., Sadarangani, M. & Lavoie, P. M. Maternal Immunological Adaptation During Normal Pregnancy. Front. Immunol. 11, (2020).

46. Landman, A. J. E. M. C. et al. Evaluation of low-dose aspirin in the prevention of recurrent spontaneous preterm labour (the APRIL study): A multicentre, randomised, double-blinded, placebo-controlled trial. PLoS Med 19, e1003892 (2022).

47. Silver, R. M. et al. Low-Dose Aspirin and Preterm Birth: A Randomized Controlled Trial. Obstetrics & Gynecology 125, 876–884 (2015).

48. Tanigawa, T. et al. Lansoprazole, a Proton Pump Inhibitor, Suppresses Production of Tumor Necrosis Factor-α and Interleukin-1β Induced by Lipopolysaccharide and Helicobacter Pylori Bacterial Components in Human Monocytic Cells via Inhibition of Activation of Nuclear Factor-κB and Extracellular Signal-Regulated Kinase. J. Clin. Biochem. Nutr. 45, 86–92 (2009).

49. Gilman-Sachs, A. et al. Inflammation induced preterm labor and birth. Journal of Reproductive Immunology 129, 53–58 (2018).

50. US Preventive Services Task Force. Folic Acid Supplementation to Prevent Neural Tube Defects: US Preventive Services Task Force Reaffirmation Recommendation Statement. JAMA 330, 454–459 (2023).

51. Wu, Y. et al. The association between periconceptional folic acid supplementation and the risk of preterm birth: a population-based retrospective cohort study of 200,000 women in China. Eur J Nutr 60, 2181–2192 (2021).

52. Armstrong, C. ACOG Guidelines on Psychiatric Medication Use During Pregnancy and Lactation. afp 78, 772–778 (2008).

53. Wang, J. et al. Timing of Antidepressant Use in Pregnancy and Preterm Birth: A Systematic Review and Meta-analysis. O&G Open 1, 022 (2024).

54. Deenick, E. K., Pelham, S. J., Kane, A. & Ma, C. S. Signal Transducer and Activator of Transcription 3 Control of Human T and B Cell Responses. Front. Immunol. 9, 168 (2018).

55. Durant, L. et al. Diverse Targets of the Transcription Factor STAT3 Contribute to T Cell Pathogenicity and Homeostasis. Immunity 32, 605–615 (2010).

56. Côté, F. et al. A novel modulator of IL-6R prevents inflammation-induced preterm birth and improves newborn outcome. EMBO Molecular Medicine 1–33 (2025) doi:10.1038/s44321-025-00257-9.

57. Wang, L. et al. IL-37 Exerts Anti-Inflammatory Effects in Fetal Membranes of Spontaneous Preterm Birth via the NF-κB and IL-6/STAT3 Signaling Pathway. Mediators Inflamm 2020, 1069563 (2020).

58. Marzioni, D. et al. Importance of STAT3 signaling in preeclampsia (Review). Int J Mol Med 55, 58 (2025).

59. Passerini, L. et al. STAT5-signaling cytokines regulate the expression of FOXP3 in CD4+CD25+ regulatory T cells and CD4+CD25− effector T cells. International Immunology 20, 421–431 (2008).

60. Burchill, M. A., Yang, J., Vogtenhuber, C., Blazar, B. R. & Farrar, M. A. IL-2 Receptor β-Dependent STAT5 Activation Is Required for the Development of Foxp3+ Regulatory T Cells. The Journal of Immunology 178, 280–290 (2007).

61. Gomez-Lopez, N., StLouis, D., Lehr, M. A., Sanchez-Rodriguez, E. N. & Arenas-Hernandez, M. Immune cells in term and preterm labor. Cell Mol Immunol 11, 571–581 (2014).

62. Aneman, I. et al. Mechanisms of Key Innate Immune Cells in Early-and Late-Onset Preeclampsia. Front. Immunol. 11, 1864 (2020).

63. Bert, S., Ward, E. J. & Nadkarni, S. Neutrophils in pregnancy: New insights into innate and adaptive immune regulation. Immunology 164, 665–676 (2021).

64. Weichhart, T. et al. The TSC-mTOR Signaling Pathway Regulates the Innate Inflammatory Response. Immunity 29, 565–577 (2008).

65. Weichhart, T., Hengstschläger, M. & Linke, M. Regulation of innate immune cell function by mTOR. Nat Rev Immunol 15, 599–614 (2015).

66. Tong, A., Huang, J., Wolf, G., Dijk, D. V. & Krishnaswamy, S. TrajectoryNet: A Dynamic Optimal Transport Network for Modeling Cellular Dynamics. in Proceedings of the 37th International Conference on Machine Learning 9526–9536 (PMLR, 2020).

67. Zhang, S., Afanassiev, A., Greenstreet, L., Matsumoto, T. & Schiebinger, G. Optimal transport analysis reveals trajectories in steady-state systems. PLoS Comput Biol 17, e1009466 (2021).

68. Bunne, C., Papaxanthos, L., Krause, A. & Cuturi, M. Proximal Optimal Transport Modeling of Population Dynamics. in Proceedings of The 25th International Conference on Artificial Intelligence and Statistics 6511–6528 (PMLR, 2022).

69. Ryu, J., Bunne, C., Pinello, L., Regev, A. & Lopez, R. Cross-modality Matching and Prediction of Perturbation Responses with Labeled Gromov-Wasserstein Optimal Transport. Preprint at 10.48550/ARXIV.2405.00838 (2024).

70. Demir, A., et al. sc-OTGM: Single-Cell Perturbation Modeling by Solving Optimal Mass Transport on the Manifold of Gaussian Mixtures. Preprint at 10.48550/ARXIV.2405.03726 (2024).

71. Dong, M. et al. Causal identification of single-cell experimental perturbation effects with CINEMA-OT. Nat Methods 20, 1769–1779 (2023).

72. Zunder, E. R. et al. Palladium-based mass tag cell barcoding with a doublet-filtering scheme and single-cell deconvolution algorithm. Nat Protoc 10, 316–333 (2015).

73. Behbehani, G. K. et al. Transient partial permeabilization with saponin enables cellular barcoding prior to surface marker staining. Cytometry Pt A 85, 1011–1019 (2014).

