## Supplemental Materials for "Single-cell-level digital twins for preterm birth prevention strategies"

### Supplementary data

**Figure S1** Infographic depicting the SIMCO workflow in detail.
**Figure S2** Unified gating strategy for mass cytometry of the perturbation dataset and the clinical dataset by Stelzer et al. 2021.
**Figure S3** Design of the 96-well barcoding scheme.
**Figure S4** Overview heatmaps of drug effects achieved by all drug candidates.
**Figure S5** Synergistic, additive, and antagonistic effects of SA, lansoprazole, and SALPZ.
**Figure S6** Modeling of time-dependent effects in signaling response across columns.

**Table S1** Antibody panel of the perturbation atlas mass cytometry experiments.
**Table S2** CellOT performance in cross-validation across stimulation conditions.
**Table S3** Predictive performances for simulated data by SIMCO, original data by Stelzer et al. 2021, and EGA.
**Table S4** List of hyperparameter grid search for Stabl/XGBoost models.

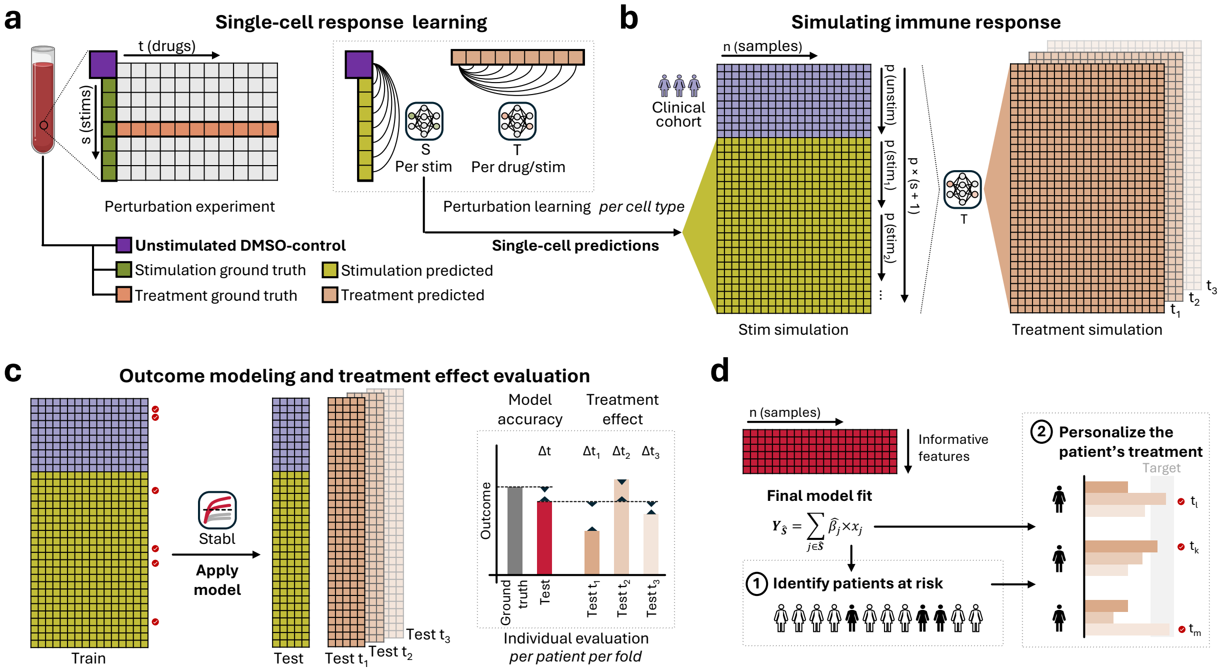

**Figure S1 | Infographic depicting the SIMCO workflow in detail.**

**a.** From a whole blood sample, perturbation experiments are performed to generate ground truth single-cell data (e.g., mass cytometry) over a matrix of s stimulations and t drug treatments.

**b.** Per cell type, CellOT models $S_v$ and $T_k$ are trained to learn perturbation responses to each stimulation v and to each treatment k (within a specific stimulation condition).

**c.** From an independent clinical cohort, n baseline samples (i.e., unstimulated and unexposed to treatments) are analyzed with mass cytometry, providing p features per sample. The resulting baseline dataset is expanded to include v stimulation scenarios through CellOT single-cell predictions, resulting in a simulated dataset containing (v +1) x p features. Using the stimulation-specific $T_k$ CellOT models, a final dataset is built simulating single-cell treatment responses for each stimulation condition. The simulated treatment-response dataset contains (v +1) x p x t features.

**d.** The Stabl algorithm is then used to build a multivariable model predictive of a given clinical outcome (e.g., the time to labor onset). The expanded dataset (before simulated treatment) is split into a training and testing cohorts for cross-validation. Stabl selects clinical outcome-informative features and builds a multivariable model on the training cohort. The resulting Stabl model is then applied to the testing cohort and its treatment-simulated datasets $t_k$.

**e.** Model accuracy is evaluated as the difference between true and predicted outcome on the testing cohort. Treatment effects are evaluated per patient per cross-validation fold as the difference between the predicted outcome on the untreated data and the treatment-stimulated data.

**f.** The final Stabl model can be used to identify patients at risk for a given clinical outcome, and evaluation on the simulated-treatment responses can help personalize the patient's treatment.

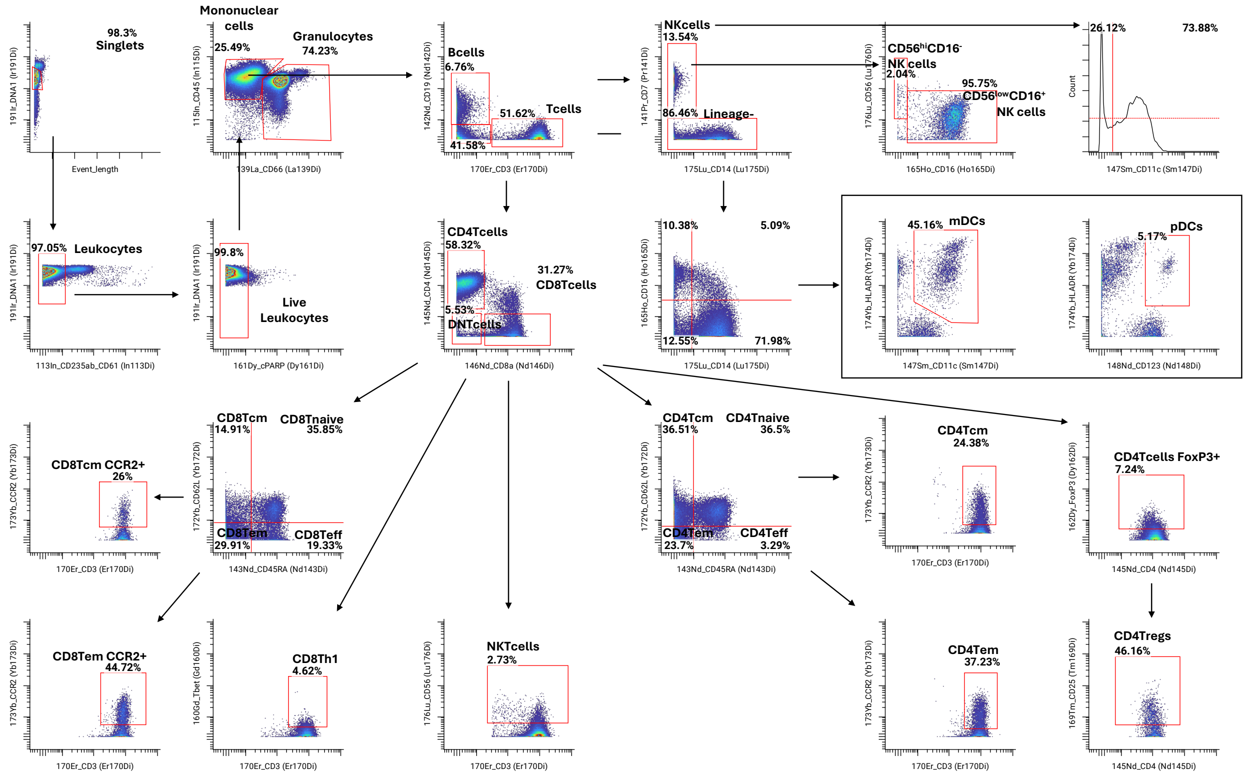

**Figure S2 | Unified gating strategy for mass cytometry of the perturbation dataset and the clinical dataset by Stelzer et al. 2021.** A total of 28 cell populations was manually gated and used for the analysis.

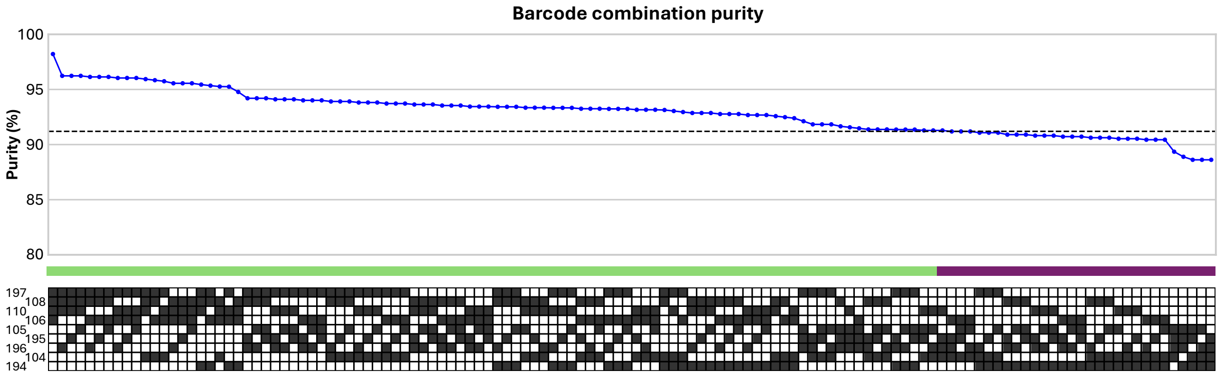

**b**

| Isotope | Purity |
|---|---|
| Au197 | 99.9 |
| Pd108 | 99.5 |
| Pd110 | 99.4 |
| Pd106 | 99.3 |
| Pd105 | 98.8 |
| Pt195 | 97.3 |
| Pt196 | 97.3 |
| Pd104 | 97.0 |
| Pt194 | 96.5 |

**c**

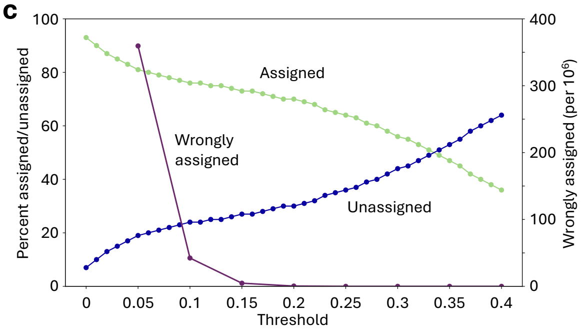

**Figure S3 | Design of the 96-well barcoding scheme.** Palladium (104, 105, 106, 108, 110), platinum (194, 195, 196) and gold (197) isotopes were used to construct a nine-choose-four barcoding system.
**a.** The theoretical purity of each of 126 possible barcoding labels calculated as product of the individual isotopes' purities. The 96 purest barcodes (green) were used, while the 30 most impure barcodes (purple) served as control. The threshold between used and unused barcodes was a purity of 91.28 %.
**b.** Table of isotope purities based on the vendor's specifications.
**c.** Varying the debarcoding threshold demonstrated how lower thresholds achieve higher assignment rates, while assigning more cells to unused ("wrong") barcodes (purple).

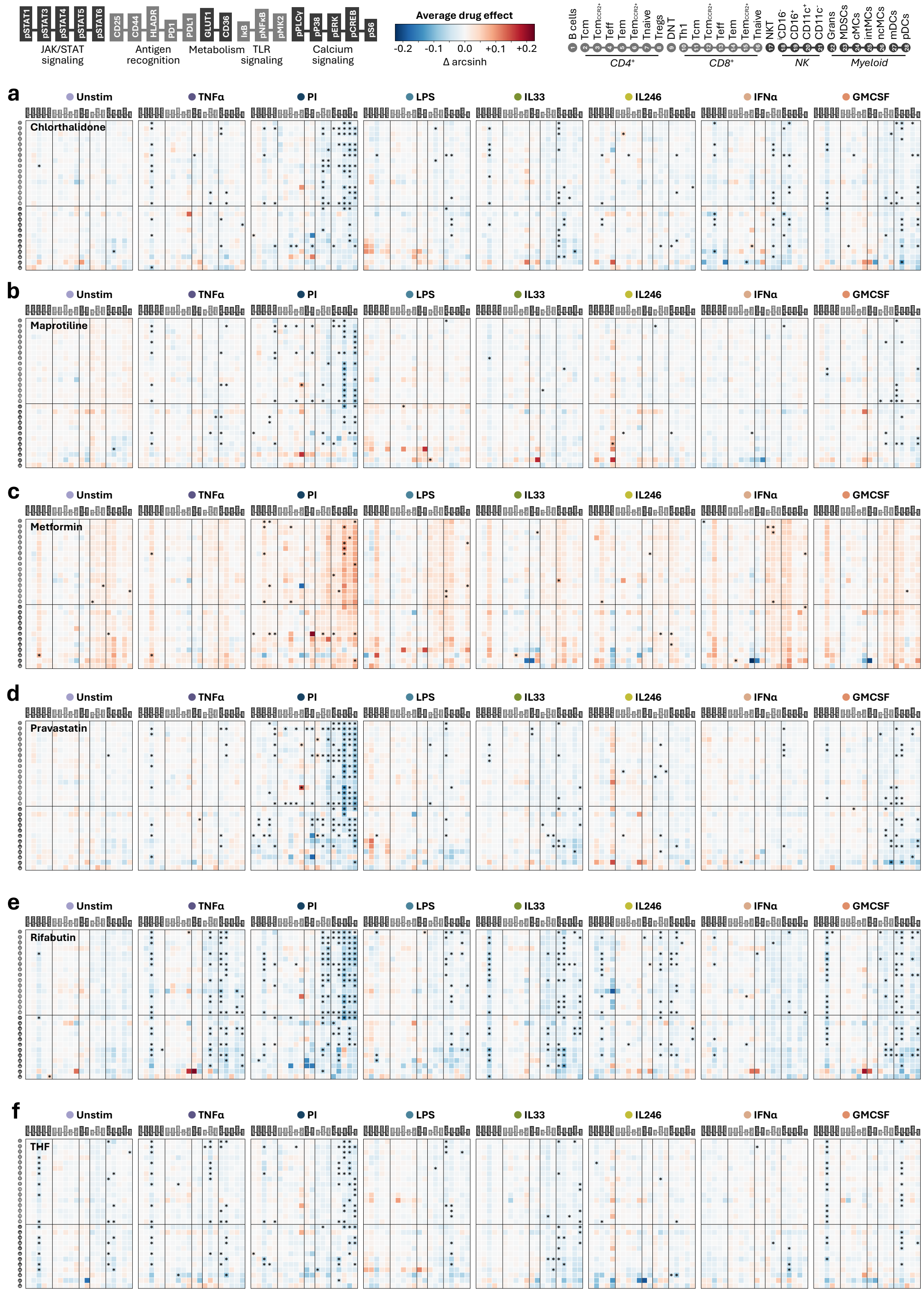

**Figure S4 | Overview heatmaps of drug effects achieved by all drug candidates.** Average drug effects across n = 11 pregnant individuals for chlorthalidone (**a**), maprotiline (**b**), metformin (**c**), pravastatin (**d**), rifabutin (**e**), and 5-methyl-tetrahydrofolate (THF, **f**). For SA, SALPZ, and lansoprazole see **Figure S5**.

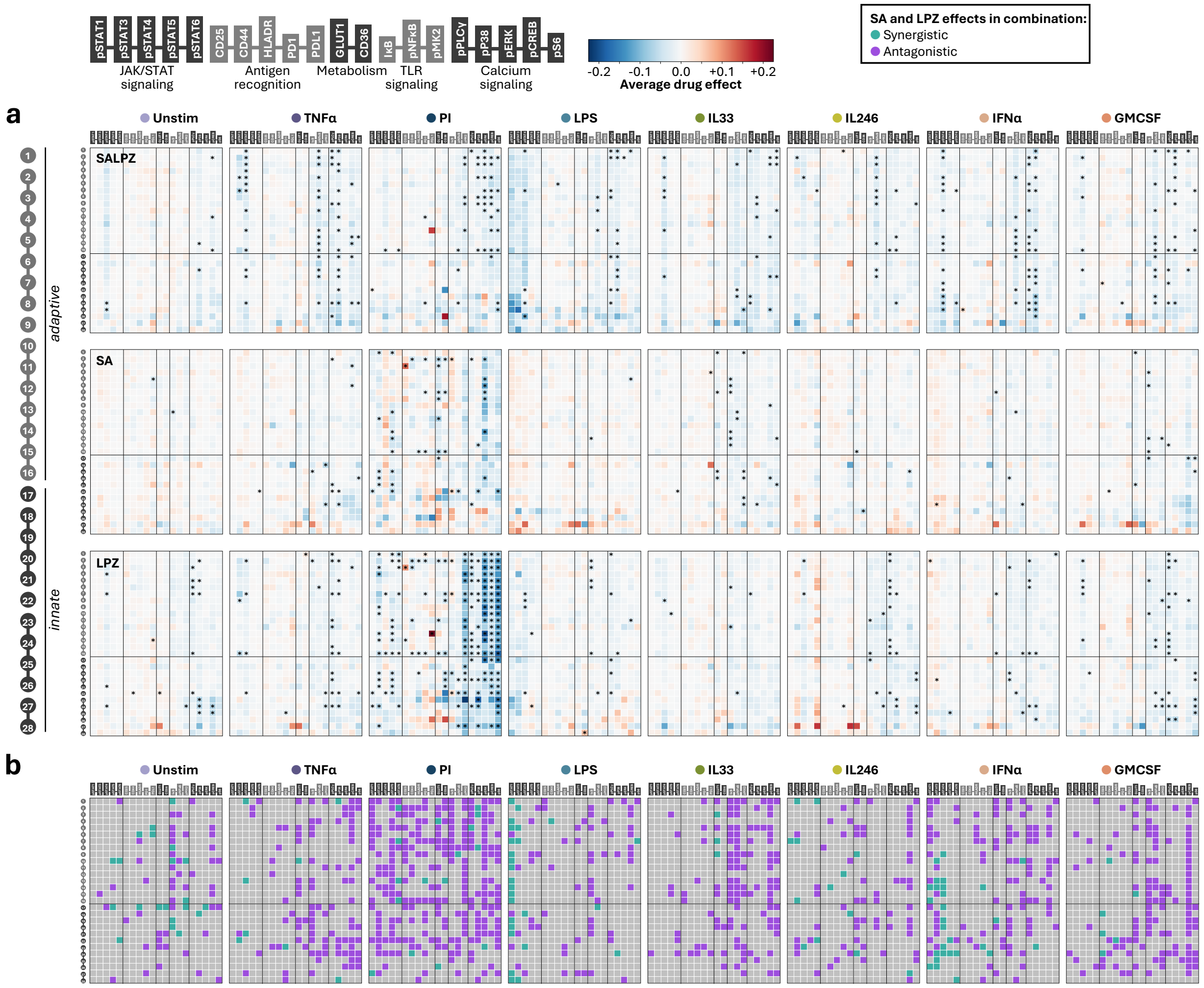

**Figure S5 | Synergistic, additive, and antagonistic effects of SA, lansoprazole, and SALPZ.**

**a.** Average drug effects mediated by SALPZ (top), SA (middle), and lansoprazole (LPZ, bottom) on n = 11 pregnant individuals.

**b.** Comparing the observed effects of SALPZ to the summed effects of SA and lansoprazole classified the effects as synergistic, additive, or antagonistic. Across all features, SALPZ effects were synergistic in 3%, additive in 79%, and antagonistic in 18%.

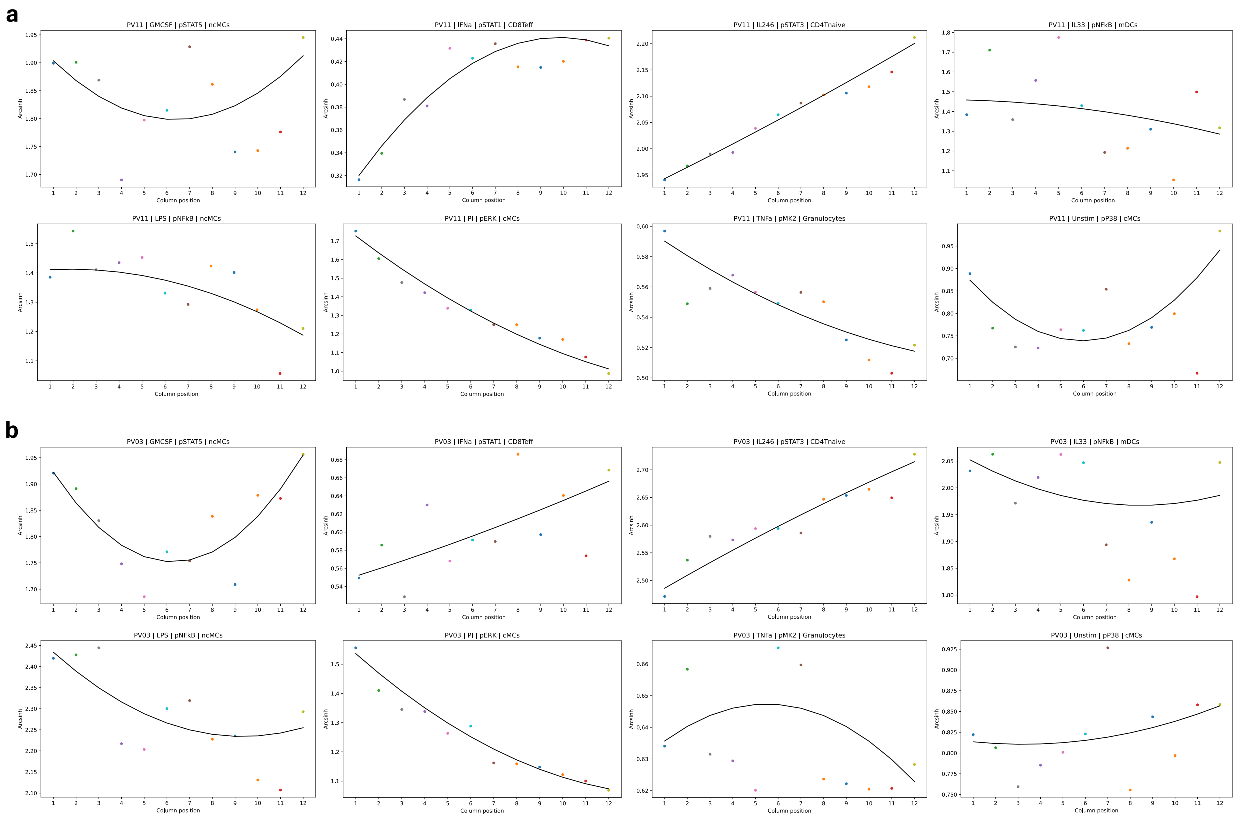

**Figure S6 | Modeling of time-dependent effects on signaling response across columns.** For individuals PV11 (**a**) and PV03 (**b**), the modeled time-dependent effects on signaling responses across columns is shown for eight example features. Per individual and per feature, a smooth spline was fitted across the feature values of all columns (k = 2, smoothing = 1.5, weight = 1 for drug-columns, weight = 2 for DMSO columns) to account for time-dependent effects in signaling response. Drug treatments were organized as depicted in Figure 2a. DMSO columns were placed in column 1, 6, and 12.

| Target | Clone | Supplier | Catalog # |
|---|---|---|---|
| CD235ab | HIR2 | Biolegend | 306615 |
| CD61 | VI-PL2 | BD | 555752 |
| CD45 | HI30 | Biolegend | 304045 |
| CD66 | CD66a-B1.1 | BD | 551354 |
| CD7 | M-T701 | BD | 555359 |
| CD19 | HIB19 | Biolegend | 302247 |
| CD45RA | HI100 | Biolegend | 304143 |
| CD11b | ICRF44 | Biolegend | 301337 |
| CD4 | RPA-T4 | Biolegend | 300541 |
| CD8a | RPA-T8 | BD | 557084 |
| CD11c | Bu15 | Biolegend | 337221 |
| CD123 | 6H6 | Biolegend | 306027 |
| PD-1 | D4W2J | CST | 63815 |
| CD44 | IM-7 | Biolegend | 103065 |
| CD36 | D8L9T | CST | 39914 |
| PD-L1 | E1L3N | CST | 85164 |
| GLUT1 | EPR3915 | Abcam | ab252403 |
| CD16 | 3G8 | Biolegend | 302051 |
| CD25 | M-A251 | Biolegend | 356102 |
| CD3 | UCHT1 | Biolegend | 300443 |
| CD62L | DREG.200 | Thermo Fisher | BMS1015 |
| CCR2 | K036C2 | Biolegend | 357202 |
| HLA-DR | L243 | Biolegend | 307651 |
| CD14 | M5E2 | Biolegend | 301843 |
| CD56 | NCAM16.2 | BD | 559043 |
| pCREB | 87G3 | CST | 9198 |
| pSTAT5 | C11C5 | CST | 51879SF |
| pp38 | 36/p38 | BD | 612281 |
| pSTAT1 | 14/P-STAT1 | BD | 612132 |
| pSTAT3 | M9C6 | CST | 74309SF |
| pS6 | D57.2.2E | CST | 4858 |
| pMAPKAPK2 | 27B7 | CST | 3007 |
| Tbet | 4B10 | Thermo Fisher | 14-5825-82 |
| cPARP | F21-852 | BD | 552597 |
| FoxP3 | PCH101 | Thermo Fisher | 14-4776-82 |
| IκB | L35A5 | CST | 4814 |
| pNFκB | K10-895.12.50 | BD | 558393 |
| pERK1/2 | D13.14.4E | CST | 45899SF |
| pSTAT6 | A15137E | Biolegend | 686002 |
| pPLCγ2 | A17025A | Biolegend | 612402 |
| pSTAT4 | 38/P-STAT4 | BD | 612739 |

**Table S1 | Antibody panel of the perturbation atlas mass cytometry experiments.**

| Stimulation | Average MAE | Average MMD | Average Spearman R |
|---|---|---|---|
| GMCSF | 0.07732349 | 0.01001739 | 0.81835463 |
| IFNα | 0.08200975 | 0.01234101 | 0.78866706 |
| IL246 | 0.08949932 | 0.01130042 | 0.8079325 |
| IL33 | 0.03352717 | 0.00247631 | 0.93522383 |
| LPS | 0.1246932 | 0.02581156 | 0.57759379 |
| PI | 0.14132662 | 0.02632193 | 0.72555276 |
| TNFα | 0.04773449 | 0.00462354 | 0.90259611 |

**Table S2 | CellOT performance in cross-validation across stimulation conditions.**

| Model | AUROC | Pearson R | R2 | Global RMSE | RMSE term | RMSE preterm |
|---|---|---|---|---|---|---|
| Simulated stim features (SIMCO) + Unstim features (Stelzer) | 0.729 | 0.64 | 0.42 | 24.12 | 23.98 | 25.4 |
| Simulated stim features (SIMCO) + Unstim features (Stelzer) + EGA | 0.857 | 0.95 | 0.9 | 10.16 | 8.27 | 20.7 |
| EGA | 0.5 | 0.94 | 0.83 | 12.83 | 6.85 | 35.68 |
| Stim features (Stelzer) + Unstim features (Stelzer) | 0.788 | 0.75 | 0.55 | 21.14 | 21.57 | 16.4 |
| Unstim features (Stelzer) | 0.695 | 0.4 | 0.15 | 29.16 | 29.48 | 25.9 |
| Simulated stim features (SIMCO) | 0.715 | 0.64 | 0.4 | 24.38 | 24.2 | 25.95 |
| Simulated stim features (SIMCO) + Stim features (Stelzer) + Unstim features (Stelzer) | 0.805 | 0.8 | 0.62 | 19.41 | 19.68 | 16.5 |
| Simulated stim features (SIMCO) + Stim features (Stelzer) + Unstim features (Stelzer) + EGA | 0.9 | 0.95 | 0.9 | 9.96 | 8.72 | 17.8 |

**Table S3 | Predictive performances for simulated data by SIMCO, original data by Stelzer et al. 2021, and EGA.**

| **Stabl grid search** | |
|---|---|
| **Hyperparameters** | **Values** |
| Base estimator | [Lasso, Alasso, EN] |
| Final model fitted | [Linear regression, Random Forest, XGBoost] |
| Artificial noise | [Knockoff, Random permutation] |
| Noise proportion | [0.8, 1.0] |
| **XGBoost grid search (final layer)** | |
| **Hyperparameters** | **Values** |
| n_estimators | [300, 500] |
| max_depth | [2, 4, 10] |
| learning_rate | [0.01, 0.05] |
| subsample | [0.5, 0.7] |
| colsample_bytree | [0.5,0.8] |
| gamma | [0, 1] |
| reg_alpha | [0] |
| reg_lambda | [1] |

**Table S4 | List of hyperparameter grid search for Stabl/XGBoost models.**
